# Compartment-specific tuning of hippocampal dendritic feature selectivity by intracellular Ca^2+^ release

**DOI:** 10.1101/2021.09.27.460811

**Authors:** Justin K. O’Hare, Yusuke Hirabayashi, Victoria L. Hewitt, Heike Blockus, Miklos Szoboszlay, Sebi V. Rolotti, Tristan C. Geiller, Adrian Negrean, Vikas Chelur, Attila Losonczy, Franck Polleux

**Affiliations:** Department of Neuroscience, Columbia University; New York, NY, 10027, United States; Mortimer B. Zuckerman Mind Brain Behavior Institute, Columbia University; New York, NY, 10027, United States; Kavli Institute for Brain Science, Columbia University; New York, NY, 10027, United States; Department of Chemistry and Biotechnology, School of Engineering, The University of Tokyo; Tokyo, Japan

## Abstract

Dendritic Ca^2+^ signaling is central to neural plasticity mechanisms allowing animals to adapt to the environment. Intracellular Ca^2+^ release (ICR) from endoplasmic reticulum has long been thought to shape these mechanisms. However, ICR has not been investigated in mammalian neurons *in vivo*. We combined electroporation of single CA1 pyramidal neurons, simultaneous imaging of dendritic and somatic activity during spatial navigation, optogenetic place field induction, and acute genetic augmentation of ICR cytosolic impact to reveal that ICR supports the establishment of dendritic feature selectivity and shapes integrative properties determining output-level receptive fields. This role for ICR was more prominent in apical than in basal dendrites. Thus, ICR cooperates with circuit-level architecture *in vivo* to promote the emergence of behaviorally-relevant plasticity in a compartment-specific manner.

## Main Text

Learning occurs when experience-driven neuronal activity patterns induce changes in synaptic strengths, thereby altering how future information propagates through neuronal circuits. This process, known as synaptic plasticity, is a fundamental way in which animals adapt to the environment and yet it remains enigmatic. At the cellular level, Ca^2+^ plays an essential role in transducing specific patterns of synaptic activation into long-lasting changes in synaptic efficacy^1^. The magnitude and spatiotemporal patterning of cytosolic Ca^2+^ are therefore critical in determining which synapses will undergo plasticity and the extent to which they will do so. Most studies of the mechanisms regulating dendritic Ca^2+^ dynamics have focused on the role of voltage-gated channels mediating fast Ca^2+^ conductances at the plasma membrane^2–4^. However, another critical regulator of intracellular Ca^2+^ dynamics is represented by the main internal Ca^2+^ store, the endoplasmic reticulum (ER) network^5^, which pervades the dendritic arbor^6–8^ where it sequesters nearly all Ca^2+^ within a neuron. The ER is able to release its highly-concentrated store of Ca^2+^ in an activity-dependent manner, significantly amplifying the impact of Ca^2+^ influx from the extracellular space^5, 9–14^. Thus, the ER is poised to control the magnitude and spatial extent of synaptic plasticity by dramatically increasing local cytosolic Ca^2+^ concentration in dendritic segments receiving high degrees of synaptic input.

While significant attention has been paid to the potential role for ICR in plasticity through numerous *in vitro* studies^11–21^, and recently in *Drosophila*^22^, the potential of this powerful intracellular amplificatory process remains unexplored in mammalian neurons *in vivo*. This is due to a lack of tools to precisely and effectively manipulate ICR in awake, behaving vertebrate animal models. Such an approach would require not only cellular and pre- vs. postsynaptic specificity but also the ability to influence ICR across each of its two canonical release mechanisms^5, 23^. To overcome this limitation, we focused on *Pdzd8*; a gene recently shown to encode a tethering protein that brings ER and mitochondria into direct contact, thereby enabling mitochondria to buffer a substantial fraction of ER-released Ca^2+^ ^9^. Reduction of *Pdzd8* expression allows a greater amount of Ca^2+^ to escape into the cytosol upon synaptically-driven ICR in dendrites while leaving functional and morphological properties of ER and mitochondria intact^9^. Here we leverage *Pdzd8* as a gain-of-function tool to augment the cytosolic impact of ICR by, for the first time, deleting a gene in single neurons in adult mice *in vivo*. We deploy this novel molecular-genetic tool in pyramidal neurons of hippocampal area CA1 (CA1PNs), using ‘place fields’ (PFs) as a model system to test the role of ICR in synaptic plasticity mechanisms supporting dendritic and cellular feature selectivity. PFs are spatial receptive fields that arise in an experience-dependent manner to collectively form a comprehensive map of an animal’s environment^24, 25^ and are thus widely studied as a substrate for hippocampal learning and memory. We report that ICR promotes the emergence of dendritic feature selectivity during spatial navigation. This subcellular phenomenon cooperates with compartment-specific organization of synaptic inputs across the dendritic arbor, preferentially acting on apical versus basal dendrites, to ultimately determine output-level properties of spatial receptive fields *in vivo*.

## Single-cell genetic deletion and *in vivo* subcellular imaging in hippocampal area CA1

To understand the role of ICR in dendritic and output-level feature selectivity, we sought to simultaneously monitor activity of dendrites and their cognate somata in CA1PNs with either normal or augmented ICR during spatial navigation. Using an *in vivo* single-cell electroporation approach adapted to hippocampal area CA1 (see Methods), we introduced pCAG-Cre, pCAG-FLEX-jGCaMP7b, and pCaMKII-bReaChes-mRuby3 to single CA1PNs in anesthetized adult mice (Fig. 1A, left and middle). We targeted single cells in a new *Pdzd8* conditional knockout mouse line (*Pdzd8^F/F^*) (Fig. S1, A and B) and in *WT* control mice to generate single *Pdzd8 KO* (augmented ICR) and *WT* cells (Fig. 1A, right) expressing a high-baseline Ca^2+^ indicator (jGCaMP7b) and a fluorescently-tagged excitatory opsin (bReaChes-mRuby3). The presence of Cre recombinase reduced PDZD8 protein expression within 7 days (Fig. S1, C and D). Our acute, single-cell genetic deletion circumvented potential non-cell-autonomous^22, 26^ and developmental^16^ effects that might have arisen from broad and constitutive perturbations and further allowed us to unambiguously assign dendrites to their parent soma. During head-fixed spatial navigation on a cue-rich, 2-meter treadmill belt^27, 28^, we simultaneously monitored activity dynamics in one dendritic focal plane, which ranged from basal dendrites in *stratum oriens* (*SO*) to distal tuft dendrites in *stratum lacunosum moleculare* (*SLM*, Fig. S2), and in a somatic plane which often contained co-planar segments of basal dendrite (Fig. 1, B and C). For motion correction (Fig. S3) and dendritic ROI registration (see Methods) purposes, we co-acquired static mRuby3 signals through a dedicated collection channel (Fig. 1B).

**Fig. 1.**
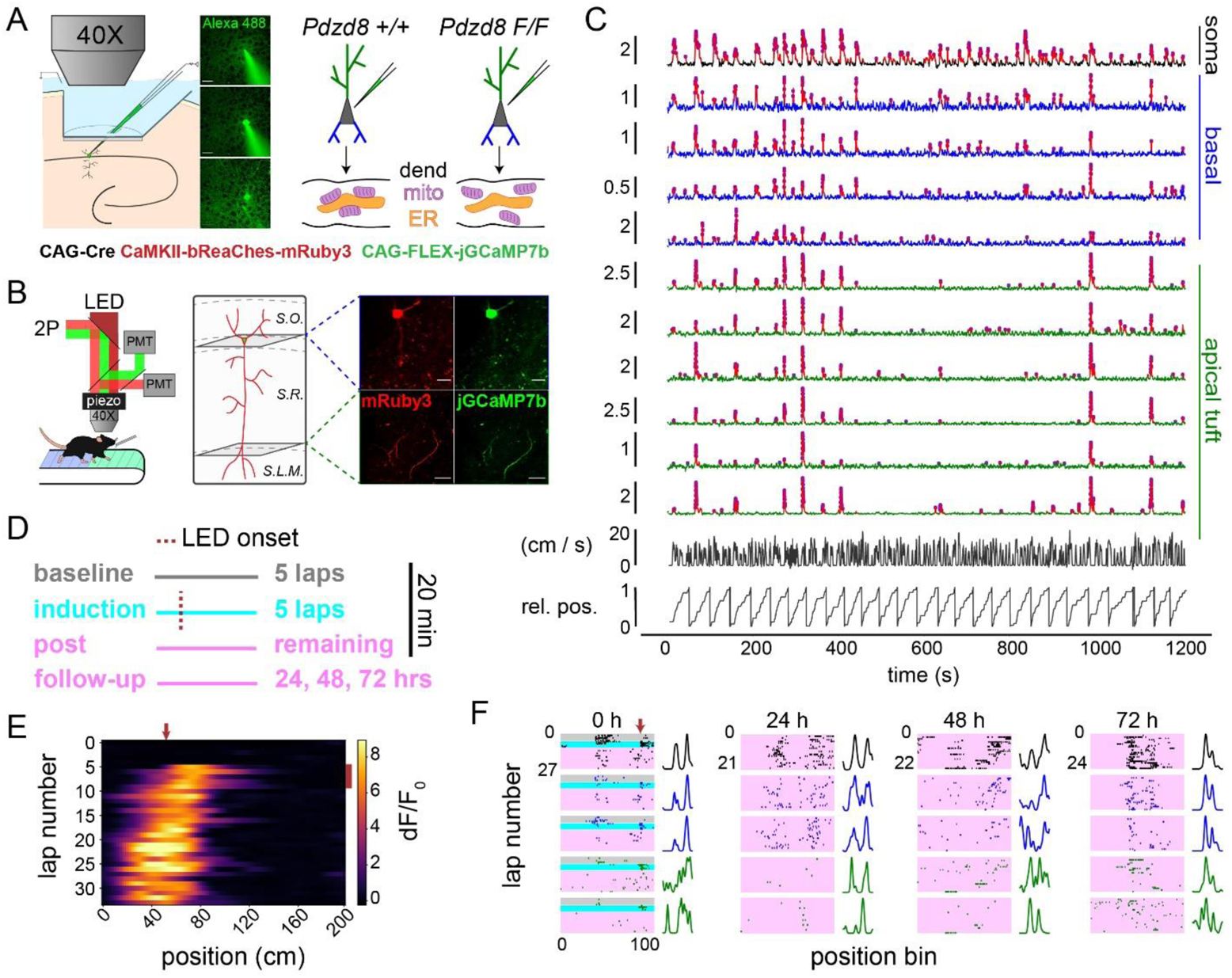
Simultaneous imaging of somatic and dendritic activity in single, spatially-tuned CA1PNs with normal and augmented ICR. (**A**) Left: An electroporation pipette containing plasmid DNAs (listed below) is guided through a silicon-protected slit in an imaging window implanted over dorsal CA1. Middle: 2-photon images before (top), during (middle), and after (bottom) electroporation of plasmid solution to a single cell in the pyramidal layer. Scale bar 20 µm. Right: Plasmid mixture is delivered via single cell electroporation to CA1PNs in *Pdzd8^+/+^* and *Pdzd8^F/F^* mice. Schematic shows Cre-mediated ablation of ER-mitochondria (mito) contacts in dendrite (dend) of *Pdzd8^F/F^* CA1PN, augmenting cytosolic impact of ICR. (**B**) Left: Multi-plane, dual-channel imaging during head-fixed spatial navigation on a spatially-cued belt. LED (620 nm) is used for photoactivation of the opsin-expressing cell. Right: Diagram approximating focal planes imaged for an example cell with corresponding motion-corrected, time-averaged fields of view from each channel showing soma/co-planar basal dendrites (top row) and distal apical tuft dendrites (bottom row). Scale bar 50 µm. Hippocampal strata indicated by dashed lines: *S.O., stratum oriens; S.R., stratum radiatum; S.L.M., stratum lacunosum moleculare*. (**C**) Example traces of soma (black), as well as basal (blue) and apical tuft (green) dendrites co-acquired from example cell in (B). Vertical scale bars indicate dF/F_0_. Detected Ca^2+^ transients drawn in red with deconvolved events as magenta circles. Frame-synchronized animal running speed and relative position (rel. pos.) shown below traces. (**D**) Schematic describing optogenetic place field induction paradigm. (**E**) Example somatic activity heatmap from induction session. LED location denoted by red arrow at top; LED laps by red bar at right. (**F**) Example event rasters and spatial tuning curves from a single CA1PN showing somatic and subset of dendritic dynamics on induction day (0 h) and at 24 h, 48 h, and 72 h timepoints. ROI types colored as in (C); conditions colored as in (D). Tuning curves are scaled to maximum values to aid visualization.

To evaluate the role of ICR in plasticity and learning, we imaged CA1PNs expressing PFs. To boost the fraction^27, 29^ of CA1PNs functioning as ‘place cells’, and to track the stability of a subset of PFs with a defined time-zero, we carried out an optogenetic PF induction protocol^30^ in each imaged cell (Fig. 1, D and E). Optogenetically-induced PFs displayed characteristic hallmarks (Fig. S4, A-D) of naturally-occurring *in vivo* place cell formation^30–32^ and shared similar properties with spontaneous, non-induced PFs (Fig. S4, E-G). We therefore pooled spontaneous and induced PFs except in longitudinal analyses addressing the stability of spatial feature selectivity, where a defined time point for PF formation was necessary. We imaged somatic and dendritic activity dynamics in 20-minute sessions on four consecutive days (Fig. 1F, Fig. S5) including the initial PF induction day, comparing post-induction dynamics in CA1PNs with normal and augmented ICR.

## ICR controls diversity and reliability of dendritic spatial tuning in single CA1PNs

Given the absence of *in vivo* functional imaging data for apical CA1PN dendrites and the paucity of such data for basal dendrites^33–35^, we first sought to (1) characterize spontaneous Ca^2+^ transient properties in apical and basal dendrites with respect to their parent soma and (2) assess any impact of our ICR-augmenting manipulation on basic CA1PN activity dynamics. *Pdzd8 WT* and *KO* CA1PN cell bodies did not differ in Ca^2+^ transient frequency or amplitude (Fig. S6, A and B). As previously reported in *WT* basal dendrites^33, 35^, isolated dendritic transients (no coincident somatic event, see Fig. S6, C and D for examples) were overall rare compared to somatic transients with minor but significant differences in event frequency between compartment and genotype (Fig. S6E). Isolated apical dendritic Ca^2+^ transients, which have not previously been reported in CA1PNs, were larger and more frequent than those observed in basal dendrites (Fig. S6, E and F) while increasing ICR via *Pdzd8 KO* did not affect isolated transient amplitude in either dendritic compartment (Fig. S6F). Overall, these results are in agreement with extensive *in vitro* evidence suggesting that dendritic ICR requires a high degree of temporally correlated pre- and postsynaptic activation^5, 9, 11, 12^ and therefore is likely to play a privileged role in synaptic plasticity as opposed to regulating cytosolic Ca^2+^ dynamics associated with baseline neuronal activity patterns. Thus, *Pdzd8* appears to be a suitable lever to assess the *in vivo* impact of ICR on synaptic plasticity underlying feature selectivity in soma and dendrites.

We reasoned that, if ICR regulates synaptic plasticity *in vivo*, then augmenting its spread and magnitude should impact specific aspects of dendritic feature selectivity in CA1PNs. As with other gain-of-function manipulations, such as artificially activating specific neuronal populations^36, 37^, these effects would in turn provide insight into the endogenous function of ICR. We thus surveyed spatial tuning properties of apical and basal dendrites, with respect to their cognate soma, in individual *Pdzd8 WT* and *KO* CA1PNs. Since optogenetically-induced PFs represented only a fraction of those observed across all four days (Fig. 2, A and B), and induced PFs shared similar characteristics with spontaneous PFs (Fig. S4E-G), we initially analyzed PFs regardless of their location relative to prior LED stimulation. In *Pdzd8 KO* cells with augmented ICR, both apical and basal dendritic activity showed increased overall spatial tuning (Fig. 2C, see Methods). *Pdzd8 KO* apical and basal ‘place dendrites’, i.e. dendrites in which PFs were detected, fired more reliably upon the animal’s traversal of their PFs (Fig. 2D) while basal place dendrites fired more selectively within their PFs (Fig. 2E). Increasing the cytosolic impact of ICR did not affect any measure of somatic spatial tuning (Fig. 2, C-E). This is consistent with the previous finding that increased within-field basal dendritic activity does not portend increased CA1PN somatic activity^33^ but may also be due to relatively low sample sizes for somatic ROIs; all three metrics in Fig. 2, C-E trended higher for *Pdzd8 KO* CA1PN somata. The feature selectivity-promoting effects of increased ICR on CA1PN apical and basal dendrites, particularly given that the same manipulation does not affect isolated Ca^2+^ transient amplitude (Fig. S5F), indicates that ICR indeed regulates synaptic plasticity underlying hippocampal PF properties *in vivo*.

**Fig. 2.**
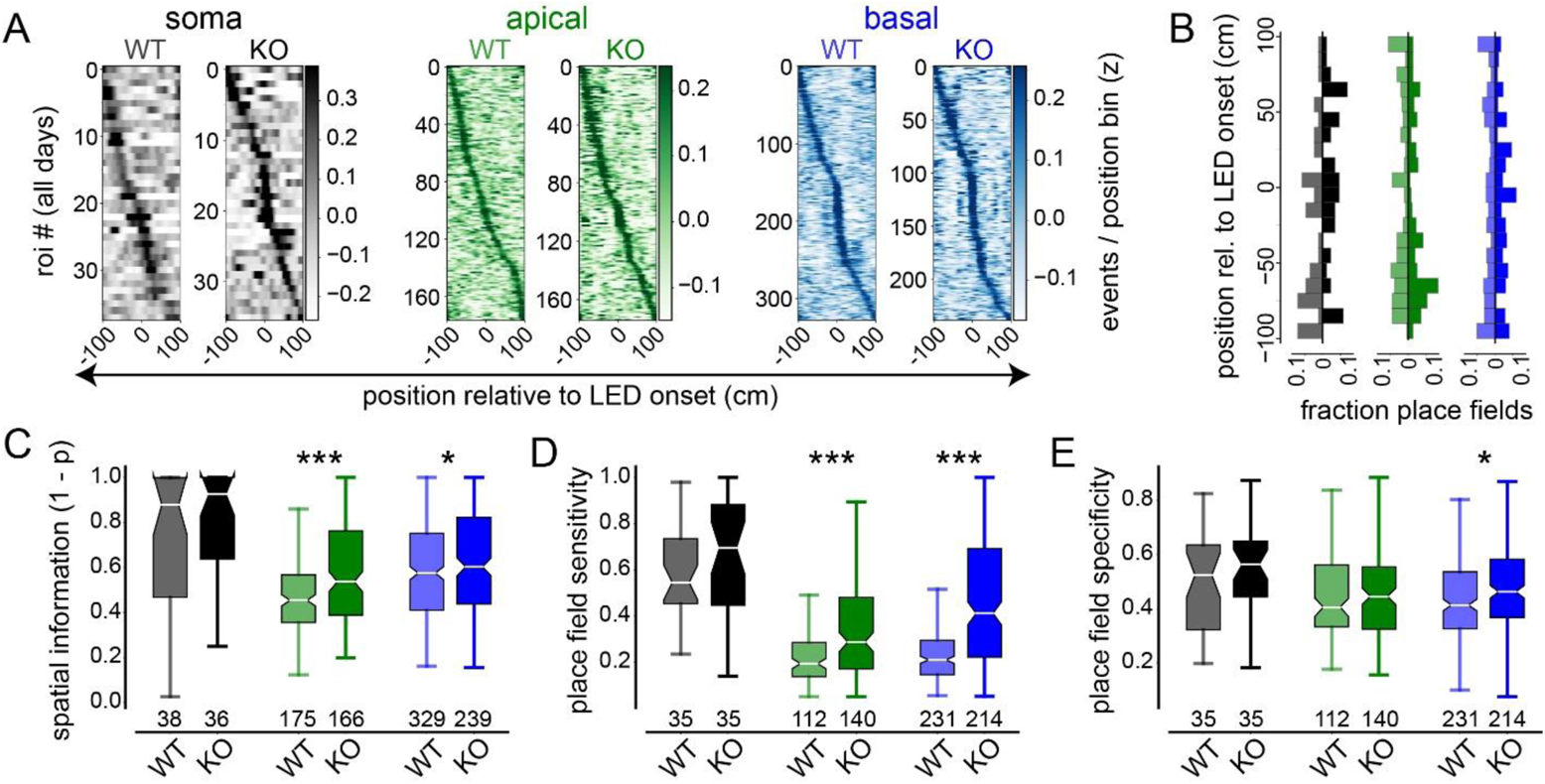
Effect of augmented ICR on somatic and dendritic spatial tuning properties in single CA1PNs. (**A**) Spatial activity heatmaps showing LED-centered spatial tuning curves for all *Pdzd8 WT* and *KO* somatic (greys), apical (greens), and basal (blues) ROIs imaged across all days, sorted by tuning curve peak location. (**B**) Mirrored vertical histograms showing distribution of place field (PF) center locations relative to LED onset for *Pdzd8 WT* (transparent; left of x-axis) and *KO* (opaque; right of x-axis) for somatic, apical, and basal ROIs color-coded as in (A). (**C-E**) PF metrics, including spatial information (C), PF sensitivity (D), and PF specificity (E), shown for all ROI type-genotype combinations. Boxes range from lower to upper quartiles with line at median; whiskers show range of data within 1.5 * (Q3 - Q1) of box boundaries. For panels (A-C), N represents total number of ROIs imaged across days (see Table S1 and Methods for details). For panels (D, E), N represents total number of ROIs with significant spatial tuning across days. Two-sided unpaired t-tests and Mann-Whitney U tests were used. p < 0.05*, 0.001***.

We next tested whether ICR influences the distribution of dendritic spatial tuning preferences (Fig. 3A). Augmenting ICR by *Pdzd8 KO* significantly increased the correlation of spatial tuning curves (TCs) among connected apical dendritic branches (Fig. 3B, see Fig. S7 for illustration of analytic approach) as well as between apical dendrites and their soma (Fig. 3C). TC correlations were curiously unaffected in basal dendrites, although we note that *WT* basal dendrites already displayed high dendrite-dendrite and dendrite-soma correlations which may have led to a ceiling effect (Fig. 3, B and C). To exclude dendrites that exhibited minimal spatial tuning to begin with, we next restricted our analysis to place dendrites that belonged to place cells and calculated the minimum circular distances along the cued belt that separated each dendritic PF from that of its soma. This analysis revealed a previously unappreciated degree of heterogeneity in spatial tuning preference among *WT* apical dendrites relative to their somata and, in contrast, a clear tendency toward somatic co-tuning in *WT* basal dendrites (Fig. 3D). Moreover, increased ICR was associated with a significant shift in apical dendritic PFs toward those of their soma; *Pdzd8 KO* apical dendrites were as co-tuned to their soma as were *WT* basal dendrites (Fig. 3D). Consistent with the observation that *Pdzd8 KO* selectively affected apical dendrite TC correlations (Fig. 3, B and C), *Pdzd8 KO* did not affect basal dendrite-soma co-tuning (Fig. 3D). Notably, dendritic PF specificity did not attenuate with increasing soma-dendrite PF distance as might be expected if anti-tuned dendritic PFs were not *bona fide* receptive fields (Fig. S8). In fact, apical PFs in *Pdzd8 KO* CA1PNs displayed increasing specificity the more anti-tuned they were with their somatic PFs (Fig. S8A, right). Additionally, neither branch order nor path length predicted dendrite-soma co-tuning (Fig. 3E). This is consistent with the prior observation that co-firing between basal dendrites and their cell bodies does not depend on intervening distance^33^ and indicates that somatic action potential backpropagation does not appreciably influence soma-dendrite co-tuning, at least across the ranges of branch order and path length that we sampled (Fig. S2). Finally, we formally tested the extent of dendrite-soma co-tuning by training a regularized linear model to predict somatic TCs based on those of connected dendrites (Fig. 3F, see Methods). This model revealed that apical dendrites with augmented ICR markedly outperformed *WT* apical dendrites in predicting their somatic TCs while the same manipulation caused basal dendrites to marginally underperform (Fig. 3G). Taken together, these results indicate that ICR promotes reliable dendritic feature selectivity while specifically enhancing co-tuning of apical dendrites, with each other and with their soma, across the apical arbor.

**Fig. 3.**
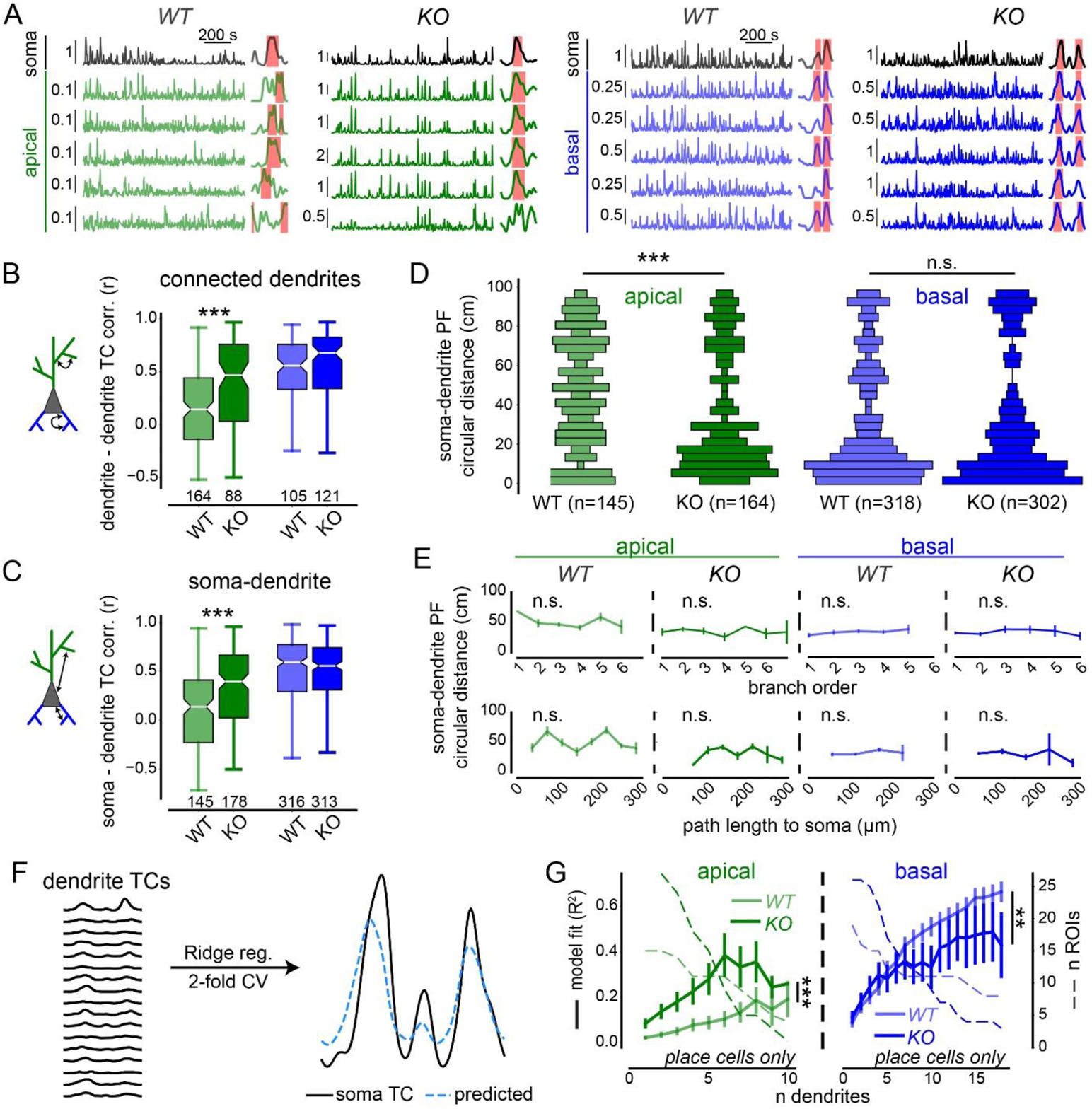
Compartment-specific regulation of dendrite-dendrite and soma-dendrite co-tuning by ICR. (**A**) Example dF/F_0_ traces and tuning curves (TCs) for all ROI type-genotype combinations. Vertical scale bars indicate dF/F_0_. Dendritic traces and TCs are grouped under corresponding somata. Detected PFs are denoted by pink shaded areas overlaid onto TCs. (**B-C**) TC correlations calculated exclusively between connected, co-imaged dendrites (B, see Fig. S7) and between dendritic ROIs and their corresponding somata (C). Ns represent total number of ROIs imaged across days (Table S1). Boxes range from lower to upper quartiles with line at median; whiskers show range of data within 1.5 * (Q3 - Q1) of box boundaries. (**D**) Vertical histograms showing distributions of minimum circular distances between somatic and dendritic place field (PF) centers. Bar widths represent relative abundance of bin values. Ns represent total PFs from imaged dendrites belonging to place cells across days. (**E**) Circular distances in (D) plotted against branch order (top) and path length to soma binned every 40 µm (bottom). Spearman correlation analysis was used on non-binned data. (**F**) Schematic for predicting somatic TC based on TCs of connected dendrites (see Methods) with example performance shown at right. (**G**) Model performance (R^2^), plotted against the number of dendrites included in training data. Number of cell-sessions containing N dendrites shown in dashed lines on second y-axis. Note, measurement precision decreases with N. Apical 2-way ANOVA, genotype effect: F_1,43_ = 37.39, p < 0.001); n dendrites effect: F_9,217_ = 4.57, p < 0.001; interaction: F_9,217_ = 0.81, p > 0.05; n = 16 *WT* and 28 *KO* place cells. Basal 2-way ANOVA, genotype effect: F_1,44_ = 7.35, p < 0.01; n dendrites effect: F_17,387_ = 13.44, p < 0.001; interaction: F_17,387_ = 1.05, p > 0.05; n = 19 *WT* and 26 *KO* place cells. Error bars represent SEM. Distributions in (B-D) were compared using two-sided unpaired t-tests and Mann-Whitney U tests. p < 0.01**, 0.001***.

## ICR differentially shapes the subcellular distribution of spatial tuning within single apical and basal dendritic segments

The amount of Ca^2+^ released from ER into the cytosol influences CA1PN dendritic spatial tuning *in vivo*, but how does the extent of ICR fit into plasticity mechanisms underlying hippocampal feature selectivity? Presence of ER in dendritic spines positively predicts spine head size^8^, suggesting a role for ICR in spine structural plasticity. *In vitro* experiments support such a role^38–40^ and further suggest that the spread of ICR along a dendritic branch in part determines which spines will undergo synaptic plasticity^41, 42^. Therefore, the spatial extent of ICR along a dendritic branch may ultimately influence local integrative properties underlying dendritic spatial receptive fields *in vivo*. To investigate this possibility, we next asked how increasing the cytosolic impact of ICR would influence activity and feature selectivity within single dendrites, i.e. across nearby dendritic spines. We subdivided dendritic ROIs into 2-micron segments (“subROIs”) and re-extracted, processed, and analyzed subROI signals (Fig. 4A). To account for within-dendrite differences in focality, we normalized activity measures to static mRuby3 signal intensity (see Methods). We found that variability in overall activity levels between subROIs was selectively reduced in *Pdzd8 KO* apical dendrites (Fig. 4B). Next, we asked whether increasing ICR alters spatial features of isolated Ca^2+^ transients along individual dendritic segments (Fig. 3C, Fig. S9A). Indeed, genetic amplification of ICR increased the spread of isolated Ca^2+^ transients specifically in apical dendrites (Fig. 4D, Fig. S9, B-E, see Methods). Finally, we calculated TC correlations between all possible combinations of subROIs and asked how co-tuning varied between subROIs as a function of the anatomical distance separating them along the dendritic arbor (Fig. 4E). Both apical and basal *WT* dendrites showed a monotonic decrease in intradendritic TC correlation with distance within single dendritic branches (Fig. 4F). Given the relatively low sensitivity of genetically-encoded Ca^2+^ indicators such as jGCaMP7b^43^, this effect indicates that feature-correlated inputs are spatially clustered along CA1PN dendrites as has previously been suggested^16, 44–47^ and recently demonstrated^48^. Strikingly, increasing ICR strengthened intradendritic TC correlations in both compartments and erased the distance-dependent drop-off in TC correlations specifically within apical dendrites (Fig. 4F). Together, these results identify a role for ICR in shaping the distribution of synaptically-driven feature selectivity along single apical and basal dendrites.

**Fig. 4.**
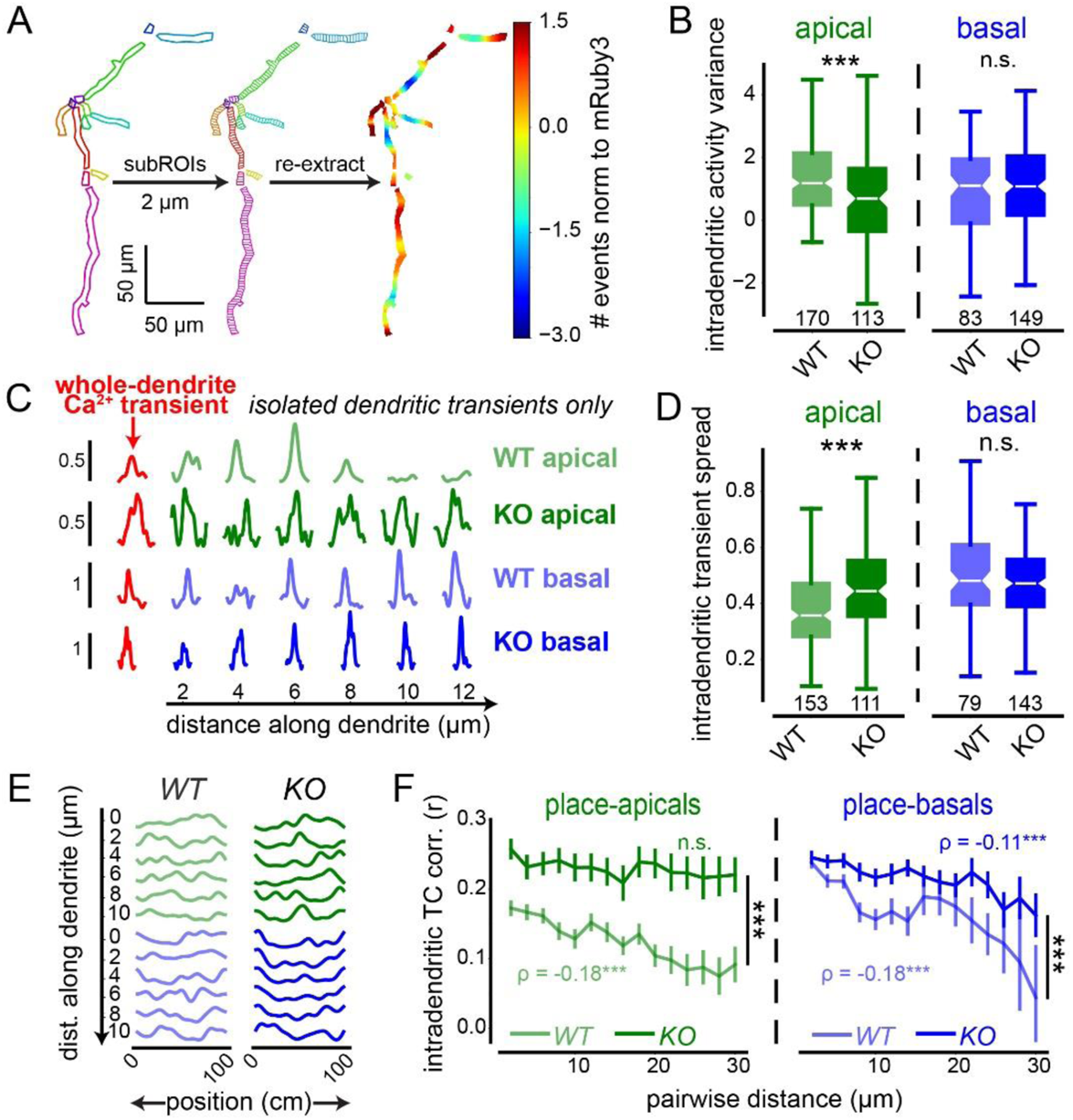
Intracellular Ca^2+^ release shapes activity dynamics and spatial feature selectivity within single dendritic branches of CA1 PNs. (**A**) Approach to analyze intradendritic dynamics. Each ROI is segmented into 2-μm subROIs along its longest axis. Signals are then re-extracted from each subROI for further analysis. Heatmap shows activity distribution for each imaged apical dendrite from a single example *WT* CA1PN. Activity is normalized within-ROI to static mRuby3 signal intensity (log_2_ ratio) to control for differences in focality (see Methods). (**B**) Within-dendrite variance of activity levels between *WT* and *Pdzd8 KO* apical and basal dendrites (log_2_ ratio of coefficients of variation for events and mRuby3 signal intensity). (**C**) Spatial spread of isolated Ca^2+^ transients within single dendrites. For each Ca^2+^ transient detected from a whole-dendrite ROI (red traces), corresponding dF/F_0_ signals are plotted from the first 6 subROIs (12 μm) of that dendrite. (**D**) Normalized intradendritic spread of isolated Ca^2+^ transients within *WT* and *Pdzd8 KO* apical and basal dendrites. A value of 1.0 indicates uniform subROI peak amplitudes (see Methods). (**E**) Example subROI spatial tuning curves (TCs), color-coded as in (C). (**F**) Intradendritic TC correlations plotted as a function of distance separating two subROIs (see Methods). Spearman correlation coefficients are shown on plots. Distances were compared between genotypes using Mann-Whitney U tests. N = 110 *WT* and 100 *KO* place-apicals; 223 *WT* and 184 *KO* place-basals. Error bars represent SEM. Boxes range from lower to upper quartiles with line at median; whiskers show range of data within 1.5 * (Q3 - Q1) of box boundaries. Distributions in (B and D) were compared using two-sided unpaired t-tests. ***p < 0.001.

## ICR strengthens dendritic feature selectivity and stabilizes output-level receptive fields

CA1 place cells can gain, lose, remap, or retain their spatial tuning properties. Exposure to a new environment promotes PF formation and remapping^49, 50^ while, upon repeated exposure to the same environment, subsets of cells stably represent specific locations^29, 51, 52^. This balance between dynamism and stability is thought to enable animals to flexibly adapt to new environments while remembering familiar environments. Given the role of ICR in controlling the spatial extent of feature selectivity within and across dendrites, and in promoting reliability in dendritic PFs, we tested how these input-level PF properties ultimately influence output-level, i.e. somatic, spatial tuning over days. To assess the relative stability of CA1PN spatial tuning over days, we restricted our analysis to activity dynamics within a nominal ‘LED zone’ centered around the location of LED onset during our optogenetic induction protocol (Fig. 5, A-C). We longitudinally tracked somatic activity in the same environment across 72 hours and assessed dendritic activity specifically on day 0 (see Methods). Since PF formation requires both pre- and postsynaptic activity and only a subset of synapses should receive excitatory presynaptic input at the time of postsynaptic optogenetic stimulation, we expected to induce a fraction of dendrites. Still, dendrites of CA1PNs with augmented ICR were more likely to display optogenetically-induced spatial tuning (Fig. S10, A-C), revealing a role for ICR in the establishment of dendritic feature selectivity. To measure the strength of induced spatial tuning, we focused specifically on ROIs that were successfully induced (see Methods). We first accounted for cell-to-cell differences in plasmid expression, excitability, and/or focality by normalizing dF/F_0_ from each ROI to its peak LED response. We then quantified induced PF strength based on post-induction activity near the location of LED onset relative to a baseline measurement (Fig. 5C).

**Fig. 5.**
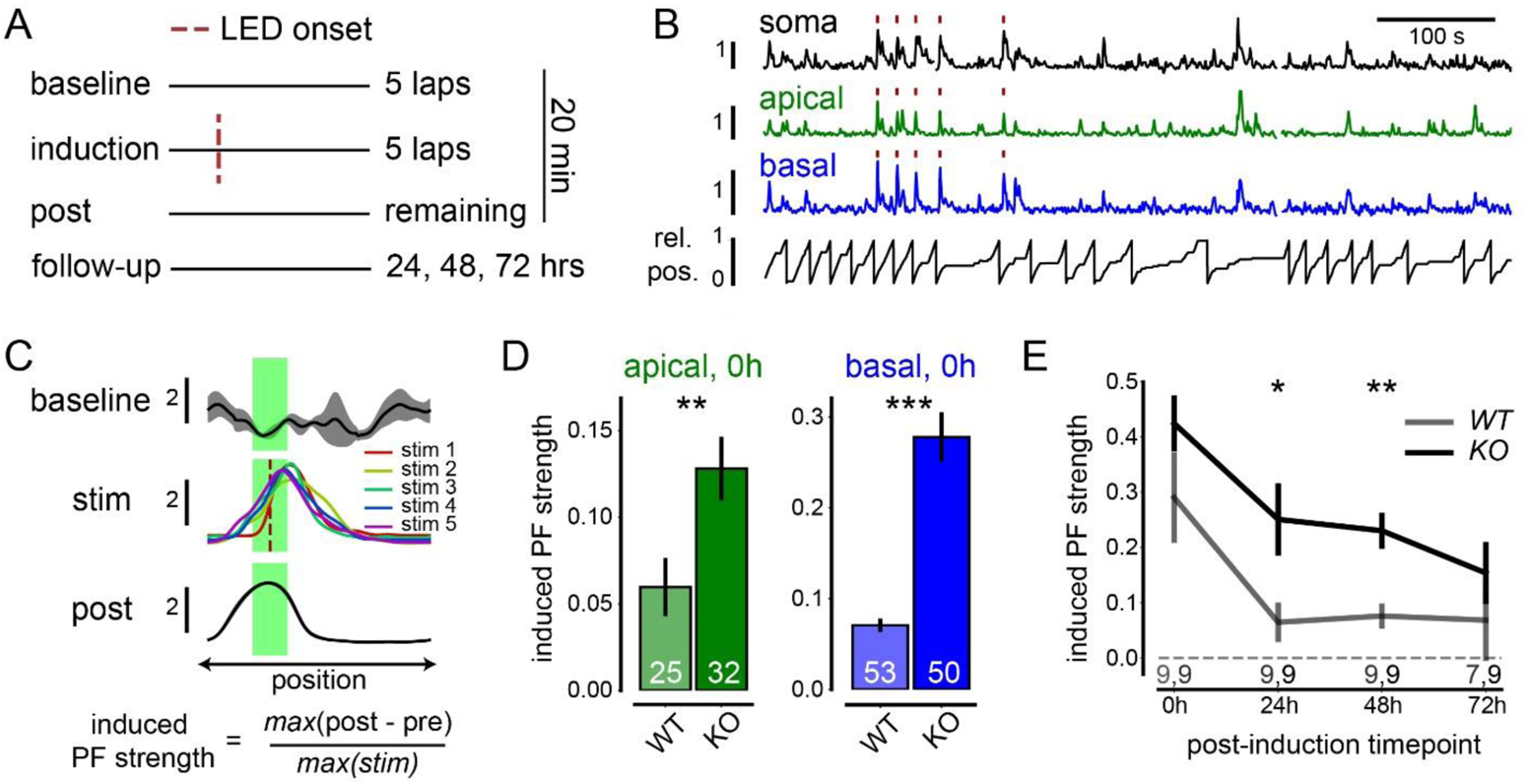
Dendritic strength and somatic stability of optogenetically-induced place fields in single *WT* and *Pdzd8 KO* CA1PNs. (**A**) Paradigm for optogenetic place field (PF) induction as in Fig. 1D. (**B**) Example traces showing LED-evoked and naturally-occurring activity in simultaneously acquired somatic, apical, and basal jGCaMP7b traces from a single CA1PN with relative animal position (rel. pos.) plotted below. LED stimulation during induction laps indicated by red ticks. (**C**) Quantifying strength of induced spatial tuning. Top: Example mean somatic dF/F_0_ by position from baseline, induction (stim), and post laps for an induced cell with baseline and post activity shown as mean ± SEM and induction laps shown individually. Red dashed line indicates LED onset and green shaded area denotes “LED zone” used for quantifying induced activity. Bottom: Quantification of induced PF strength. Signals are normalized to max LED response to control for cell-to-cell variability in jGCaMP7b expression level, excitability, and/or focality. (**D**) Apical and basal induced PF strength for successfully induced dendrites of *WT* and *Pdzd8 KO* CA1PNs on day 0. Two-sided unpaired t-tests were used. (**E**) Somatic induced PF strength of successfully induced CA1PNs across days. Mixed effects model with post-hoc *t* tests, genotype effect: F_1,16_ = 7.80, p < 0.05; time effect: F_3,46_ = 10.85, p < 0.001; interaction: F_3,46_ = 0.32, p > 0.05. p < 0.05*, 0.01**, 0.001***. Error bars and shaded error bands indicate SEM.

Both apical and basal dendrites of *Pdzd8 KO* CA1PNs displayed stronger induced PFs on Day 0 (induction day, Fig. 5D). While *Pdzd8 KO* cell bodies did not show significantly stronger PFs on Day 0 relative to *WT* cells, consistent with our observation that increased dendritic spatial tuning did not significantly translate to output-level PFs (Fig. 2, C-E), induced somatic PFs were more stable over days (Fig. 5E). Backpropagating somatic activity does not likely explain stronger dendritic PFs in *Pdzd8 KO* CA1PNs; somatic induced PF strengths were similar on Day 0 and dendritic proximity to soma does not predict soma-dendrite co-tuning (Fig. 3E). Therefore, increasing the cytosolic impact of ICR strengthens dendritic tuning and ultimately stabilizes output-level receptive fields.

## ICR shapes place field formation to strengthen and stabilize output-level spatial tuning

To more rigorously assess the impact of increasing ICR on output-level feature selectivity, and to do so in the context of naturally-occurring spatial tuning, we next imaged large, intermingled populations of *Pdzd8 KO* and *WT* CA1PN cell bodies (see Fig. S11D, Methods for classification strategy) as mice navigated a familiar cued treadmill belt over five consecutive days (Fig. 6, A-C). ROIs were registered across days for longitudinal monitoring of activity dynamics and spatial tuning (Fig. 6C, bottom; figs. S11 and S12). This large-scale, population-level analysis uncovered changes in somatic PF properties (Fig. 6, D-F) that were not detectable with sample sizes in single-cell experiments (Fig. 2, C-E). Additionally, imaging *Pdzd8 KO* and *WT* CA1PNs within-animal revealed that augmenting ICR widens output-level PFs (Fig. 6G); this comparison was not possible in the between-subjects design of single-cell experiments due to the relationship between velocity at time of PF formation and width of the resultant PF^31^ (figs. S4C and S10, D-F). This systematic increase in *Pdzd8 KO* somatic PF width is consistent with amplified ICR broadening the anatomical distribution of spatial tuning within dendritic segments (Fig. 4F). While *KO* CA1PCs were curiously less likely to display significant spatial tuning (Fig. 6H), those that did continued to do so more consistently (Fig. 6I) and displayed more-stable TCs over days relative to *WT* control cells (Fig. 6J) as observed in optogenetically-induced activity (Fig. 5E). We also noted that, while place cell turnover was pronounced (Fig. 6I)^29, 51^, CA1PNs that were place cells on either day *N* or day *N* - 1 showed increasingly stable TCs with repeated exposure to a familiar environment (Fig. 6J).

**Fig. 6.**
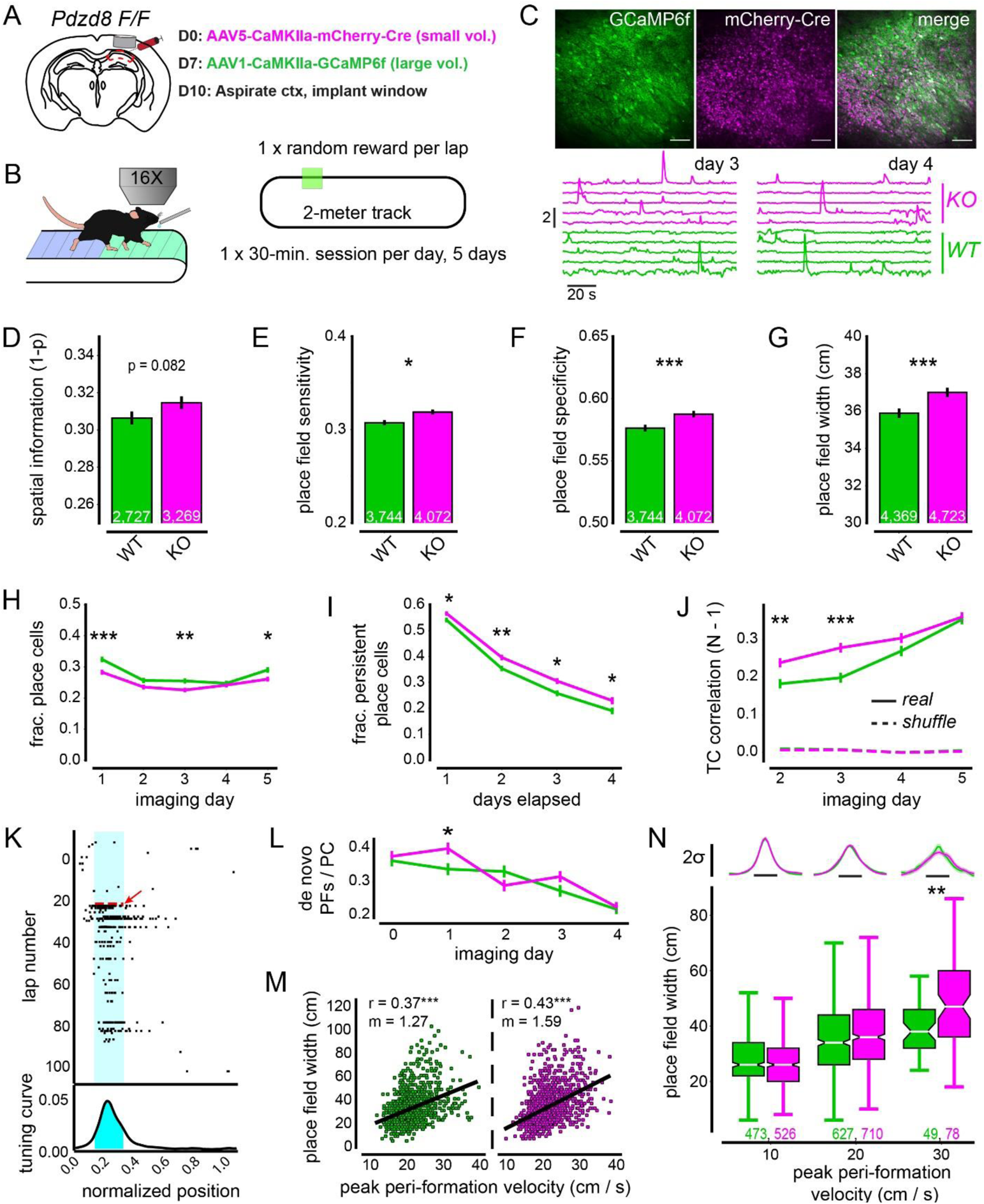
Large-scale, mixed population imaging of *WT* and *Pdzd8 KO* CA1PNs implicates intracellular Ca^2+^ release in formation and stability of output-level feature selectivity. (**A**) Viral strategy. AAV encoding mCherry-Cre fusion protein is injected days before GECI-encoding AAV at low volume to allow for recombination/protein clearance and robust mCherry signal. (**B**) Signals are acquired at 16X (1.2 - 1.5X optical zoom) during a head-fixed spatial navigation paradigm with randomly located reward. Identical fields of view (FoVs) are imaged across 5 consecutive days (see Fig. S11). N = 2,719 *WT* and 3,260 *KO* cells from 6 mice. (**C**) Top: example FoVs showing GCaMP6f (left) and mCherry-Cre (middle) expression with overlay (right). Scale bar 100 µm. Bottom: Example GCaMP6f traces from the same *Pdzd8 KO* (magenta) and *WT* (green) ROIs imaged on separate days. (**D-G**) Spatial tuning properties of *Pdzd8 KO* and *WT* cells including spatial information (D) as well as place field (PF) sensitivity (E), specificity (F), and width (G). (**H**) Fraction of place cells over days. 2-way repeated measures ANOVA with post-hoc *t* tests, genotype effect: F_1,5994_ = 10.22, p < 0.01; time effect: F_4,23976_ = 35.23, p < 0.001; interaction: F_4,23976_ = 2.09, p > 0.05 (**I**) Fraction of cells showing persistent spatial selectivity (independent of PF location) as a function of days elapsed. 2-way ANOVA with post-hoc *t* tests, genotype effect: F_1,1808_ = 23,65, p < 0.001; time effect: F_3,15959_ = 383.05, p < 0.001; interaction: F_3,15959_ = 0.51, p > 0.05. (**J**) Tuning curve (TC) correlations over days, relative to previous day. 2-way ANOVA with post-hoc *t* tests, genotype effect: F_1,3394_ = 23.86, p < 0.001; day effect: F_3,10182_ = 50.37, p < 0.001; interaction: F_3,10182_ = 2.92, p < 0.05. Dashed lines represent TC correlations calculated from shuffled data. (**K**) Example plot showing automated detection of *de novo* PF formation event. Top: deconvolved events (black ticks) plotted by lap number and position along cued belt. Red dashed line and arrow indicate detected formation lap with PF bounds shaded in cyan. Bottom: TC for above raster plot with PF shaded in cyan. (**L**) Rates of *de novo* PF formation over days for all place cells. 2-way ANOVA with post-hoc *t* tests, genotype effect: F_1,1808_ = 2.41, p > 0.05; time effect: F_4,7806_ = 28.99, p < 0.001; interaction: F_4,7806_ = 2.65, p < 0.05. (**M**) Relationship between peri-formation velocity and resultant PF width for *Pdzd8 WT* (left, green, N = 1,149) and KO (right, magenta, N = 1,314) events. Pearson correlation coefficients (r) and linear regression fit slopes (m) are shown on plots. 2-way ANOVA shows interaction between genotype and peak peri-formation velocity (F_1, 1897_ = 4.82, p < 0.001). (**N**) PF width plotted for *Pdzd8 WT* and *KO* PNs across three velocity bins. Mean TCs (z-score) plotted above for visual comparison. Horizontal scale bar 50 cm. Error bars represent SEM. Boxes range from lower to upper quartiles with line at median; whiskers show range of data within 1.5 * (Q3 - Q1) of box boundaries. Distributions in (D-G and N) were compared using two-sided unpaired t-tests and Mann-Whitney U tests. p < 0.05*, 0.01**, 0.001***.

The increased PF width observed in *Pdzd8 KO* CA1PCs (Fig. 6G) strongly suggests that ICR participates in PF formation; the synaptic plasticity rule underlying PF formation determines initial PF width^31^. To more closely examine this finding, we devised a method to automatically detect hundreds of spontaneous PF formation events in CA1PNs with normal and augmented ICR (Fig. 6K, see Methods). Spontaneous formation events occurred at similar rates between *Pdzd8 KO* and *WT* CA1PNs (Fig. 6L) and both genotypes displayed robust correlations between peri-formation velocity and PF width (Fig. 6M), supporting our detection strategy and the prevalence of behavioral time scale synaptic plasticity^31, 50^ in spontaneous PF formation. We found that augmenting ICR altered this relationship: the linear fit of *Pdzd8 KO* data was steeper (*WT:* m = 1.27, *KO*: m = 1.59) and a two-way interaction existed between genotype and peak peri-formation velocity (Fig. 6M). Binning PF width by peri-formation velocity revealed that PFs were wider in *Pdzd8 KO* CA1PNs specifically when mice ran at high speeds during PF formation (Fig. 6N). This selective effect of increased ICR on PF width at higher running speeds is consistent with a velocity-dependent increase in the frequency of position-related presynaptic CA3 input^53, 54^ arriving onto the postsynaptic CA1 cell^55^. Since CA1PN synapses require temporally-correlated pre- and postsynaptic activity to undergo plasticity *in vivo*, higher running speed increases the pool of synapses eligible for potentiation; hence the velocity-PF width relationship^31^. We therefore suggest that, while normal ICR may reach and potentiate most eligible spines at lower velocities, increased ICR more comprehensively engages larger pools of eligible connections generated during bouts of increased running speed.

## Discussion

This study represents the first functional investigation of ICR in mammalian neurons *in vivo*. Overall, we find that release of Ca^2+^ from the endoplasmic reticulum plays a key role in synaptic plasticity mechanisms underlying dendritic receptive field formation, regulating feature selectivity at multiple levels of organization; from intra- and inter-dendritic levels to dendrite-soma co-tuning. Notably, these dendritic functions coalesce to govern multiple properties of output-level spatial tuning. The combination of approaches we developed offers new insights into how ICR participates in PF formation. Based on the velocity-dependent effect of increased ICR on PF width, we conclude that this process plays a role in the formation of PFs. We can also conclude that ICR operates downstream of PF-forming dendritic plateau potentials: somatic PF stability was improved both in spontaneously-occurring as well as optogenetically-induced PFs in which plateau potentials and their associated global Ca^2+^ influx^32, 56^ are obviated by direct optogenetic depolarization. It may seem counterintuitive that the spread of ICR would be important if plateau potentials (or optogenetic stimulation) already drive global Ca^2+^ influx. It is important to note, however, that voltage-gated influx of extracellular Ca^2+^ associated even with high degrees of spiking exerts a minor impact on cytosolic Ca^2+^ concentrations ([Ca^2+^]_i_) relative to ICR^11, 14^. Moreover, computational modeling suggests that voltage-gated Ca^2+^ influx is insufficient to drive the changes in [Ca^2+^]_i_ required for associative plasticity^57^. In this light, we suggest that the spread of ICR, as regulated by ER-mitochondria contact sites^9^, determines the distribution of local feature selectivity and thus effectively transduces presynaptic input and postsynaptic bursting into lasting changes in synaptic efficacy^31, 58^. Given the recent observation that spatially-correlated presynaptic inputs from CA3 cluster onto adjacent spines in CA1PN apical dendrites^48^, the local spread of ICR should control the fraction of eligible synapses^31^ that are potentiated during PF formation. Indeed, local, within-dendrite TC correlations dropped off after roughly ~10 µm, the same radius reported for input clustering^34^, and ablation of ER-mitochondria contacts widened this radius as well as output-level spatial receptive fields; particularly at high running speeds when the number of synapses eligible for potentiation would have been highest^31^.

Our study also marks the first *in vivo* imaging of apical dendritic dynamics in CA1PNs which have received tremendous attention for their presumed central role in hippocampus-dependent learning. Independent of ICR, we found that apical dendritic tuning preferences are broadly distributed relative to somatic. Thus, similar to pyramidal neurons of various sensory cortices^59–61^, CA1PNs are able to integrate diverse dendritic tuning preferences into single, clean receptive fields. How this aspect of feature selectivity emerges remains poorly understood. Basal dendrites differed in this respect, showing strong correlation with somatic tuning. This may be due to a true biological difference between apical and basal dendrites of CA1PNs or it may be a result of more efficient backpropagation into basal dendrites which are relatively proximal to the soma. However, given that soma-dendrite co-tuning was predicted by neither branch order nor path length in either dendritic compartment, we propose that apical dendritic tuning is more decoupled from somatic tuning relative to basal and that ICR, as regulated by mitochondrial Ca^2+^ buffering, contributes to this functional compartmentalization. The compartment-specific nature of ICR’s role in plasticity intersects with the CA1PN input structure in a potentially powerful way. Given that apical and basal dendrites receive synaptic input from unique combinations of afferent circuits carrying distinct streams of information^62^, compartment-specific action by ER may allow selected features of experience to preferentially influence learning.

## Acknowledgments

The authors thank Dr. S. Fusi for productive discussion regarding intradendritic dynamics, A. Villegas for assistance piloting an immunoassay to validate conditional knockout strategy, and Drs. A. Nelson and M. Rossi for their invaluable comments on the manuscript.

## Author contributions

Conceptualization: JKO, AL, FP

Methodology: JKO, SVR, HB, MS, TG, AN, VLH, VC

Investigation: JKO, VLH Data curation: JKO, VC Formal analysis: JKO, VLH Software: JKO, SVR, VC, TG Visualization: JKO, YH, VLH

Validation: JKO, VC

Resources: YH, HB, MS, AL, FP Funding acquisition: JKO, AL, FP Project administration: AL, FP Supervision: AL, FP

Writing – original draft: JKO, AL, FP

Writing – review & editing: JKO, YH, VLH, VC, AL, FP

## Methods

### Animals

All experiments were conducted in accordance with NIH guidelines and approval of the Columbia University Institutional Animal Care and Use Committee. Animal health and welfare was supervised by a designated veterinarian. Columbia University animal facilities comply with all appropriate standards of care including cage conditions, space per animal, temperature, light, humidity, food, and water. 2-6 months old male and female *Pdzd8 F/F* and *WT* mice, fully backcrossed to C57Bl/6 genetic background, were used for all experiments. Authors are not aware of an influence of sex on the measures of interest to this study.

### Plasmids

pCAG-Cre was generated in the Polleux lab as previously described^63^. pAAV-CAG-FLEX-jGCaMP7b-WPRE was obtained from Addgene (#104497) and used as-is. pAAV-CaMKIIa-bReaChes-mRuby3 was subcloned from pAAV-CaMKIIa-bReaChes-EYFP (gift from Karl Deisseroth) by excising the EYFP sequence and replacing it with an mRuby3 fragment (cloned from pAAV-CAG-mRuby3-WPRE, Addgene #107744) using the In-Fusion cloning kit (Takara Bio). A Cre-dependent jGCaMP7b construct was chosen to confirm expression of untagged Cre.

### Generation of a *Pdzd8* conditional knock out mouse line

To generate a floxed-allele *Pdzd8* (Pdzd8^F/+^) mouse line, in which the coding sequence of exon 3 of *Pdzd8* is flanked by loxP-sites (Fig. S1A), a parental mouse line Pdzd8^tm1a(EUCOMM)Wtsi^ was obtained from the Wellcome Trust Sanger Institute. Then, a cassette encoding LacZ and Neo sequences flanked by FRT sites was removed by crossing the mice with Flp-deleter mouse line B6.129S4-Gt(ROSA)26Sor^tm1(FLP1)Dym^ (obtained from the Jackson laboratory). Pdzd8^F/+^ mice were then fully backcrossed to C57Bl/6 genetic background and homozygous Pdzd8^F/F^ mice were used as conditional knockout animals.

### Western blot quantification of *Pdzd8* knockout

Embryonic mouse cortices (E15.5) were dissected in Hank’s Balanced Salt Solution (HBSS, Gibco 14185-052) supplemented with HEPES (2.5 mM, pH 7.4, Gibco, 15630-080), 30mM D-glucose (Sigma G8769), 1mM calcium chloride (Sigma-Aldrich, C5080), 1mM magnesium sulfate (Sigma-Aldrich, M2643) and 4mM sodium bicarbonate (Gibco 25080-094) and incubated in HBSS containing papain (10 U/mL, Worthington LK003178) and 5 μg/mL DNase I (Sigma-Aldrich D5025-150KU) for 20 min at 37 °C. Samples were washed three times with HBSS and dissociated by pipetting in neurobasal media (Gibco, 21103-049) supplemented with 1x B-27 Plus Supplement (Gibco A35828-01), 1x Penicillin-Streptomycin (Gibco 15140-122), 2.5% FBS (Benchmark, Gemini 100-106), 2mM GlutaMAX (1x Gibco 35050-062) and 1x N-2 Supplement (Gibco 17502-048). Cells (500 000) were plated onto 6-well culture dishes (Corning 3516) that had been treated with 0.1 mg/mL poly-D-lysine (Gibco A3890401) for 1 h before thorough washing in sterile water.

At DIV5, half the media was replaced with neurobasal media without FBS. On DIV6 each well was treated with 7.5 µL of lentiviral preparations of pSyn-Cre or pSyn-ΔCre (negative control) (gifts from Anton Maximov). Cells were harvested at DIV7 (Day 1) or DIV14 (Day 7) in N-PER (87792 Thermo Scientific) supplemented with protease inhibitors (cOmplete^TM^ Ultra, Roche, 5892970001) and benzonase (EMD Millipore, 70664-10KUN) as per manufacturer’s instructions. Lysates were heated to 70 °C for 10 min in 1x Laemilli Sample buffer (BioRad, 1610747) supplemented with betamercaptoethanol (BioRad 161-0710). Lysates were loaded onto 4–20% Tris-Glycine gels (BioRad MiniProtean TGX, 456-1094) and run at 100 V for 15 min then 150 V for 30 min. Gels were wet transferred using Trans-blot Turbo (BioRad 1704150) High MW program onto 0.2 µm nitrocellulose (Bio-Rad 1704158). Membranes were blocked for 1 h at room temperature in Odyssey Blocking Buffer (PBS) (Li-COR, 927-4000). Primary antibodies were diluted with their respective blocking buffer with 0.1% Tween (PBS-T): PDZD8 (*9*) 1:250, Actin 1:10,000 (EMD Millipore MAB1501). Following overnight incubation at 4 °C, washes were performed with PBS-T and secondary antibodies were added at 1:20 000 for 1 h at room temperature. Actin was used as a loading control and PDZD8 quantifications were normalized to total protein stain (Revert 700, LICOR 926-11011).

Membranes were imaged wet on the Odyssey CLx imaging system using 700- and 800-nm channels and visualized using ImageStudio software version 5.2 (LI-COR Biosciences).

### Surgical procedures

All procedures were carried out under continuous isoflurane anesthesia with maintenance of body temperature. Subcutaneous administration of meloxicam and bupivacaine were used for long- and short-term analgesia, respectively. Following each procedure, mice were administered 1.0 mL PBS subcutaneously and allowed to recover in their home cages with external heat supply until recumbency was regained. Post-operative health checks were carried out twice-daily for three days following surgery, after which behavioral training could commence. Specific details by experiment follow.

#### Single-cell imaging experiments

After surgical preparation, the skull was exposed and a 3 mm craniotomy was made, centered at AP −2.2, ML −1.75 relative to Bregma. Dura was removed and cortex was slowly aspirated with continuous irrigation using cold, sterile 1X PBS until fiber tracts above the hippocampus became visible and the area was clear. A custom stainless-steel imaging cannula (InterPRO Additive Manufacturing Group, CT, US) with an angled ramp for pipet entry, fitted to a custom-etched glass coverslip (Potomac Photonics, MD, US) with a rectangular opening protected by a 0.2 mm layer of silicone film to protect the otherwise-exposed rectangular area (Fig. 1A, left), was inserted through the craniotomy and secured first by Vetbond veterinary adhesive and then by dental acrylic. A custom-machined stainless-steel headpost for head fixation was embedded in the dental acrylic resting atop intact skull caudal to the imaging cannula. The acrylic cap was shaped to retain a small volume of PBS for electrical grounding during electroporation.

#### Population imaging experiments

Stereotactic viral injections to dorsal CA1 were carried out as previously described^64^. Briefly, low-volume (16-20 nL per DV coordinate; 32-40 nL total) injections of AAV5-CaMKIIa-mCherry-Cre (UNC Vector Core) were made at AP −2.2, ML −1.75, DV −1.15, −1.05 relative to Bregma in adult *Pdzd8 F/F* mice. Surgical wounds were closed with sutures and protected with a thin coat of Vetbond veterinary adhesive. After one week to allow for recovery, robust mCherry signal (Fig. 6C), and effective deletion of the *Pdzd8* gene (Fig. S1, A and B) as well as turnover of any extant PDZD8 protein (Fig. S1, C and D), a bolus (75 nL per DV coordinate; 225 nL total) injection of AAV1-CaMKIIa-GCaMP6f (Penn Vector Core) was delivered to the same AP/ML coordinates at DV −1.2, −1.1, −1.0. Three days following the second viral injection, imaging cannulae were implanted over dorsal CA1. Implants were carried out as described in the previous subsection but using standard cylindrical stainless-steel cannulae, a non-etched glass coverslip, and no silicon protective layer.

### Single-cell electroporation

Electroporation solutions contained 50 ng / µL of each plasmid (pCAG-Cre, pAAV-CAG-FLEX-jGCaMP7b-WPRE, pAAV-CaMKIIa-bReaChes-mRuby3) in the following solution (ingredients in mM): 155 K-gluconate, 10 KCl, 10 HEPES, 4 KOH, 0.33 Alexa Fluor 488 (for visualization, see Fig. 1A, middle). Solution without Alexa Fluor 488 was set to pH 7.3 and 316 mOsm. Electroporation solution was back-loaded into a long-taper (Zeitz DMZ puller; Zeitz Instrumente Vertiebs GmbH, Germany) borosilicate glass pipet (Warner Instruments, MA, US; #G200-3). Pipet tip resistance (2.5 – 6 MΩ in open solution) was monitored with a Dagan BVC-700A amplifier (Dagan Corp, MN, US), Digidata 1550B digitizer (Molecular Devices, CA, US), and Clampex software (Molecular Devices, CA). During the electroporation procedure, the animal was head-fixed under a 2-photon microscope (described below in *2-photon Ca^2+^ imaging*) and anesthetized under a ketamine/xylazine mixture with continuous heat source. Sterile 1X PBS was used to make a grounded solution in the bowl-shaped dental acrylic cap atop the imaging implant. The pipet containing fluorescent plasmid solution was lowered at an angle using a motorized micromanipulator (Scientifica Ltd., UK) through the rectangular slit in the imaging coverslip and through the thin silicone protective layer while maintaining positive pipet pressure to avoid clogging. Once in the brain, the pipet was gradually lowered to the CA1 pyramidal layer (Fig. 1A, middle) while continuously expelling low amounts of fluorescent solution to aid visualization under 920 nm 2-photon excitation. As the pipet approached a putative CA1PN, tip resistance was used to monitor proximity. After a small deflection in resistance, and while maintaining positive pressure, a train of negative voltage pulses was delivered from a stimulus isolator (AMPI, Israel) to the amplifier headstage using a custom-made switch circuit. Successful electroporation was confirmed based on a cell filling with Alexa dye and retaining the dye after careful pipet retraction (Fig. 1A, middle). Plasmid DNA expression was checked 48 hours later. To ensure successful clearance of PDZD8 protein (Fig. S1, C and D), no imaging experiments were conducted earlier than 7 days post-electroporation for either *Pdzd8 F/F* or *WT* control mice.

### Behavior

For all experiments, mice ran along a spatially-cued, 2-meter treadmill belt under “random foraging” conditions in which water reward was delivered at a pseudorandom location once per lap. Initial delivery was non-operant and was followed by a 3-second period during which the animal could receive additional operant water reward by licking with a 50% reward rate. Mice were initially trained on uncued belts made of burlap material with frequent water reward to encourage learning; the number of reward locations per lap was gradually reduced from 10 to 1 over the course of training. Mice were water-restricted to 85-90% baseline weight to motivate learning. Mice were trained until either they ran reliably for 1 reward location per lap or until enough time had passed since electroporation/viral injection; whichever occurred latest. For single-cell-electroporated mice, at least 7 days elapsed between electroporation and imaging to allow for full turnover of PDZD8 protein after acute, Cre-mediated genetic knockout (Fig. 1A, right; Fig. S1, C and D). For population imaging experiments, GCaMP6f/mCherry expression was judged on a case-by-case basis before proceeding. Mice were habituated on the same cued belts used for experiments for 1-2 days with training sessions under the *in vivo* imaging setup prior to data acquisition. For each mouse, the belt was never changed across days within a given experiment and was calibrated to the exact same length each day using a fixed landmark of blackout tape, an infrared beam, and custom software.

### *In vivo* 2-photon Ca^2+^ imaging

All imaging experiments were carried out using a 2-photon 8 kHz resonant scanning microscope (Bruker Corp, MA, US) equipped with a Chameleon Ultra II (Coherent Inc, CA, US), tuned to 920 nm for green wavelength excitation, and a Fidelity-2 (Coherent Inc) laser fixed at 1070 nm for red excitation. Excitation pathways were separately controlled and combined at the microscope. Green and red fluorescence were separated with an emission filter set (HQ525/70m-2p, HQ607/45m-2p, 575dcxr, Chroma Technology Corporation, VT, US) and collected using channel-dedicated GaAsP (7422P-40, Hamamatsu Photonics K.K., Japan) photomultiplier tubes (PMTs). A custom dual stage preamplifier (1.4 x 105 dB, Bruker Corp) was used to amplify signals prior to digitization.

#### Single-cell experiments

Single-cell imaging experiments (20 min. each) were carried out with a 40x NIR water immersion objective lens (0.8 NA, 3.5 mm working distance, Nikon USA) coupled to a piezoelectric actuator (Bruker Corp) to allow rapid toggling between somatic and dendritic focal planes. Optical zoom was set to either 1.0X or 1.5X depending on the FoV and was accounted for in calculating microns per pixel for future morphological annotation (Fig. S2). For long-distance jumps (as much as ~300 µm), the “wait time” allowed for the objective to settle after a jump before acquiring data was increased to as much at 65 ms. While this step sacrificed a degree of temporal resolution, it was key to avoiding drastic undershooting of focal planes. Thus, multi-plane imaging experiments were acquired between 5.05 and 11.89 Hz depending on jump distance. Even at the low end of this range, Ca^2+^ traces were of high quality (Fig. 1C). To aid in day-to-day consistency of imaged planes, short ~30 second recordings were acquired, rapidly extracted, and averaged to generate non-motion-corrected estimates of target planes. This step was also particularly critical for long-distance jumps and resulted in reliable cross-day tracking of dendritic ROIs once conceived and implemented (Fig. S5, bottom row). In all experiments, pockels were adjusted separately for each focal plane. Continuous 1070 nm excitation of bReaChes-mRuby3 was used when baseline jGCaMP7b signals were judged to be too dim in the dendritic focal plane to allow for satisfactory post-hoc motion correction, as was typically the case. To allow for future ROI annotation, z-projections were generated by acquiring 128-frame averages at 1-micron steps under ketamine/xylazine anesthesia (as described for single-cell electroporation procedure) using the mRuby3 signal with pre-programmed red laser power correction with imaging depth.

#### Population imaging experiments

Population imaging experiments (30 min. each) were carried out using a 16X NIR water immersion objective (0.8 NA, 3.0 mm working distance, Nikon USA) with 1.0 – 1.5X optical zoom. Two goniometers (Edmund Optics, NJ, US) were used to align the CA1 pyramidal layer to the imaging plane by rotating the behavioral apparatus ± 10° along orthogonal axes. Goniometer settings were recorded for each animal and kept consistent across days for the purpose of cross-day ROI registration. After the 1^st^ of 5 recording sessions, the imaging FoV for a given mouse was carefully aligned to that of the previous day in X/Y/Z directions using several fiduciary landmarks. Immediately prior to data acquisition, a two-color, 128-frame-averaged image of mCherry and GCaMP6f signals was acquired during a period of animal immobility for future classification of imaged cells. Data acquisition was then carried out with 920 nm excitation.

### Single-cell place field induction

Induction sessions constituted the first imaging day of each single-cell imaging experiment. Following a baseline period of 5 laps, a 1-second LED photostimulation was triggered at a fixed location along the treadmill belt for 5 consecutive laps. In the case where pre-existing somatic spatial tuning was evident (this was visually assessed offline prior to baseline periods), LED location was set to be as far away as possible from the existing putative place field (PF). To deliver LED photostimulation while simultaneously acquiring jGCaMP7b Ca^2+^ dynamics, 620 nm light from a C-mounted ultra-fast, ultra-high-powered LED (UHP-T-620-SR, Prizmatix Ltd., Israel) was passed through a dichroic mirror allowing red light to pass into the objective lens back aperture while deflecting emitted green photons to their dedicated PMT (Fig. 1B, left). The red channel used to acquire mRuby3 signals was manually switched off during photostimulation. LED was triggered by a custom “pockel-blanking” circuit that relayed an inverted Pockel cell blanking signal which is briefly activated during Y-galvonometer flyback periods and during toggling of the piezoelectric device. This approach allowed for high-powered (30 - 40mW at sample), pulsed LED stimulation during image acquisition while protecting PMTs from potentially saturating LED photons that might otherwise be incompletely deflected from of the green collection channel given high LED power. Following optogenetic PF induction, the remainder of the 20-minute imaging session was used to acquire post-induction data. All cells were tracked for 72 hours except for two *WT* cells that died at the 72-hour mark (see Fig. 5E, table S1). Of the 16 induction experiments performed, all but 3 sessions involved one CA1PN. 2 *WT* experiments and 1 *KO* experiment involved two electroporated CA1PNs in the same field of view.

### Motion correction and cross-registration

#### Single-cell data

Motion correction was carried out separately for each individual single-cell imaging experiment using a modified version of the rigid, 2-dimensional translation approach implemented in SIMA^65^ that optionally accepts multi-frame image stacks to use as motion correction targets instead of running motion-corrected time averages. For each multiplane experiment, motion correction X-Y displacements were first calculated based only on the somatic imaging plane. Cell bodies were reliably bright and straightforward to correct. Moreover, the soma-based displacements were a close enough approximation for dendritic imaging planes that they provided a useful starting point from which to generate target images to then seed future rounds of motion correction (Fig. S3A). Target-based motion correction was carried out iteratively with decreasing maximum X-Y displacements to gradually fine-tune corrections (Fig. S3, A and B). These iterative steps were crucial because, especially for deep dendritic focal planes, individual frames were often dim and contained relatively little information (Fig. S3C). After motion correction, frame-by-frame correlations to the motion-corrected, time-averaged field of view were calculated and individual frames falling below a certain threshold were masked with NaN. Thresholds were defined based on z-score of a rolling mean of frame-by-frame correlations. Across days, dendritic focal planes could not always be perfectly replicated due to difficulties associated with long jump distances between focal planes (Fig. S5). Therefore, only cell bodies were tracked across days to monitor stability of spatial tuning properties. For all other analyses, somatic and dendritic ROIs were treated as independent observations on each day. For non-longitudinal analyses, we found this approach to be most appropriate given that individual ROIs often gained, lost, or changed spatial tuning properties across days (see Fig. 1F for example).

#### Population imaging data

Individual population imaging datasets were motion-corrected using the SIMA^65^ software package (Fig. S11, A-C). After motion correction, frame-by-frame correlations to the motion-corrected, time-averaged FoV were calculated and individual frames falling below a certain threshold were masked with NaN. Thresholds were defined based on z-score of a rolling mean of frame-by-frame correlations. For each mouse, a template dataset was chosen for cross-day alignment. This template was generally on day 2 or 3 out of 5 to maximize alignment values. Affine warp matrices were calculated between the motion-corrected, time-averaged FoV of each dataset and that of the template dataset. Prior to warp matrix calculation, images per preprocessed by clipping image percentiles to 5 and 95% boundaries and equalizing image histograms on an 8 x 8 grid. Warp matrices were learned using the Enhanced Correlation Coefficient (ECC) maximization algorithm^66^ with 5,000 iterations. These warp matrices were then applied to their respective datasets to align all imaging frames to each other (Fig. S11, D and E). Affine corrections were performed using the OpenCV software library (opencv.org).

### ROI curation

#### Single-cell data

ROIs for single-cell imaging data were hand-drawn over motion-corrected, time-averaged FoVs in FIJI. ROIs were manually annotated with compartment type (apical or basal) and a binary identification sequence that encoded their branch-wise locations on the dendritic arbor based on *in vivo* z-projections of static mRuby3 signal. In rare cases where a z-projection was not available, dendritic ROIs were annotated based on information gleaned from FoVs across days if possible or else were discarded entirely. To approximate path length to soma, reverse tree traversals were used to calculate cumulative direct 3-dimensional distances (based on X-Y pixel coordinates, microns per pixel, focal plane, and axial jump distance) between ROIs along a given branching path. This process was iterated while working outward from the soma one dendritic ROI at a time until all imaged ROIs along a given branching path had been measured. Compartment type, branch order, and path length to soma were linked to dendritic ROIs of a given cell which were in turn linked to the somatic ROI. Using this approach, dendrite spatial tuning properties could be related to those of their soma and regressed against their morphological properties (Fig. 3E).

To generate subROIs within individual dendritic ROIs, a rectangular bounding box was generated along the longest axis of the ROI polygon. The rectangular box was masked by the ROI polygon to trim its outer bound and then divided into 2-µm segments along its longest axis, again based on the microns per pixel of the imaging dataset. Each subROI inherited annotated tags from its parent ROI and the identification code was appended with an index indicating its anatomical order along the whole-dendrite ROI.

#### Population imaging data

For each mouse, ROIs were automatically detected using the Suite2P^67^ software package on concatenated, co-aligned datasets. Concatenated datasets were used because Suite2P detection primarily uses Ca^2+^ dynamics, and their decorrelation from surrounding pixels, to identify ROIs. Since activity levels of individual ROIs may fluctuate from day to day, using 5 consecutive imaging sessions of data was beneficial for identifying the maximum number of quality ROIs. Detected ROIs were then visually inspected using the software’s graphical user interface to ensure they aligned with clear, individual cell bodies, did not identify obvious interneurons, were fully encapsulated within the aligned field of view, and showed clean signals across days.

To classify ROIs as mCherry(+), i.e. *Pdzd8 KO*, or mCherry(-), i.e. *WT*, the 2-channel FoV (see *In vivo 2-photon Ca^2+^ imaging*) acquired before the template dataset (see *Motion correction and cross-registration*) was used. This 2-channel FoV was aligned to the template dataset time-averaged FoV, based on the green channel and using the affine warp procedure described above. ROI polygons were then overlaid onto the co-aligned red channel (mCherry signal) to classify each ROI as mCherry-negative or -positive (Fig. S11D). Classification labels were then distributed out to matching ROIs in the remaining 4 aligned datasets.

### Ca^2+^ signal processing and event detection

For all data, baseline fluorescence was calculated as the rolling maximum of the rolling minimum of Ca^2+^ signal over a 30 second window. This baseline was used to calculate dF/F_0_. dF/F_0_ was smoothed on a 2-frame window and detrended using a rolling median calculation over a 6-second window. Ca^2+^ transients were detected according to amplitude and duration of positive- and negative-going putative events as previously described^68^. To improve Ca^2+^ transient detection, dF/F_0_ was iteratively re-calculated 3 times while masking signals corresponding to identified Ca^2+^ transients. This process also improved dF/F_0_ calculation as the baseline became more accurate, i.e. less contaminated by transients.

For the purposes of spatial tuning analysis, Ca^2+^ transient onset locations and amplitudes were binarized into event trains using the OASIS software package. This software generates deconvolved event trains from baseline-subtracted Ca^2+^ signals^69^. To generate a preliminary event train, a given trace was deconvolved according to an AR1 model with L1 penalty and pre-computed signal decay constant of 400 ms. The median and MAD from spike-free areas of this train were used to estimate noise level and a noise threshold was then defined based on this estimate (thresholds provided below in specific sections). A new train was then computed as before with minimum event “size” (i.e. confidence level) set to this noise threshold and with a sparsity parameter (set to 7). Event “sizes” were discarded and trains were returned as binary vectors with each entry corresponding to an imaging frame.

#### Single-cell data

Signal extraction and processing was carried out exactly as described above. Ca^2+^ transients were detected with a 2.5 SD threshold for somatic ROIs and a 3.5 SD threshold for dendritic ROIs to account for decreased signal-to-noise. Likewise, for OASIS event deconvolution, a 3 MAD threshold was applied to somatic events and a 5 MAD threshold was applied to dendritic events. Based on thorough visual assessments, these thresholds were found to strike the best balance between sensitivity and specificity in detecting clear events.

LED-evoked events were excluded from all analyses except when used to normalize optogenetically-induced spatial tuning (Fig. 5, C-E). Dendritic transients were considered to be “isolated”, i.e. not co-occurring with transients in their respective somata, if their onset/offset transient boundaries did not overlap in time with any detected somatic transient. This definition was intended to be more conservative than, for example, defining a maximum time difference between transient onsets, since a particularly long-duration somatic transient may overlap with a subsequent dendritic transient despite onsets being separated substantially in time. Isolated transient properties (frequency and amplitude) were normalized to somatic measures as somatic values weakly predicted, and therefore biased to some degree, dendritic measures (amplitude, basal: r = 0.21, p < 0.05; amplitude, apical: r = 0.36, p < 0.001; frequency, basal: r = −0.23, p < 0.001; frequency, apical: r = −0.39, p < 0.01, Pearson correlation analysis of *WT* data).

#### Population imaging data

Ca^2+^ signals were extracted using the Suite2P software package^67^ from each ROI across motion-corrected, cross-registered, 5-session concatenated datasets for each mouse and then copied back to their original datasets. To avoid artifacts arising from Suite2P neuropil decontamination, such as negative or extremely large dF/F_0_ values, dF/F_0_ was calculated slightly differently. First, a baseline neuropil signal was estimated from the Suite2P neuropil output using the same filtering approach described above for baseline Ca^2+^ signal. This max-min-filtered neuropil baseline provided an offset while still allowing for subtraction of fast neuropil components. Thus, dF/F_0_ was calculated as (signal – baseline_signal_) / (baseline_signal_ + baseline_neuropil_). To compare Ca^2+^ transient properties between *Pdzd8 WT* and *KO* CA1PNs without Suite2P’s activity-based ROI segmentation (Fig. S12) which may bias detection toward active cells, signals were additionally extracted from hand-drawn ROIs using a modified version of SIMA^65^ that incorporated Fast Image Signal Separation Analysis (FISSA)^70^. FISSA-extracted dF/F_0_ was calculated using the same strategy as for Suite2P-extracted signals. Since FISSA does not include neuropil signal as an output unlike Suite2P, neuropil was calculated as the difference between non-FISSA, SIMA-extracted signal and FISSA-extracted signal. Ca^2+^ transients were detected with a 2.5 SD amplitude onset threshold. For OASIS event deconvolution, neuropil-subtracted dF was used: signal – (baseline_signal_ + baseline_neuropil_). A 3 MAD threshold was used for event inclusion.

### Spatial tuning analysis

#### Spatial tuning curves and place field detection

For all data related to spatial tuning, only running frames, i.e. imaging frames during which animal velocity was greater than or equal to 1.0 cm / s, were used. Position along the 2-meter treadmill belt was divided into 100 equal ‘position bins’. As described above, deconvolved events were utilized for spatial tuning analysis as they conveniently binarize the information contained in Ca^2+^ transient onset locations and amplitudes into discrete trains of events. To generate tuning curves, spatial tuning heatmaps were calculated such that, for lap *l* and position bin *i*,

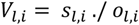

Where *s* is the total number of running frames containing deconvolved events and *o* is the total number of observations, i.e. the number of running frames falling within bin *i* on lap *l*. Spatial tuning curves were calculated as the mean of *V* along the *l*^th^ axis. Silent ROIs were excluded from spatial tuning analysis.

To determine significance of spatial tuning, 1,000 null tuning distributions were generated as above while cyclically permuting the animal position vector by a pseudorandom offset and recomputing tuning curves. Original and null tuning curves were smoothed with a 3-position bin Gaussian kernel. Significant PFs consisted of at least 5 consecutive position bins above the 95^th^ percentile of the null distribution within which an event was detected on at least 20% of laps. After PF detection, PF bounds were extended to the bin at which the spatial tuning curve intersected with:

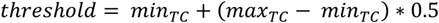

if applicable, and any overlapping PFs were merged. Dendritic spatial tuning was assessed without regard for somatic activity. For optogenetic PF induction sessions, spatial tuning analysis was carried out only using data from laps following optogenetic stimulation laps, i.e. lap 11 and onward (Fig. 1D).

#### Linear model to predict somatic tuning curves

To predict a somatic tuning curve (TC) from those of its dendrites, a linear model was trained on dendritic TCs using 2-fold cross validation, i.e. the model was given a randomly-sampled 50% of the somatic TC and was tasked with predicting the remaining 50% based on dendritic TCs. L2 regularization (Ridge regression) was used to prevent individual dendrites from being too heavily weighted. To assess model performance as a function of how many dendrites it was trained on, this procedure was iteratively run using [1:*N*] dendrites, where *N* is the total number available. Since multiple subsets could be drawn from a given set of dendrites (e.g. from 15 dendrites there are 3,003 possible combinations of 5 dendrites), the model was run 100 times per *n* and mean R^2^ values were retained. Thus, for each cell, an *N* vector of R^2^ values was returned describing model performance as a function of the number of dendrites used for training. Model predictions using all available dendrites were retained (example shown in Fig. 3F). This procedure was performed on a cell-by-cell basis and separately for each dendritic compartment.

#### Spatial information

Spatial information^71^ was chosen as a continuous measure of spatial tuning that does not rely on user-defined thresholds and parameters to identify significant PFs. This metric is agnostic of ‘place cell’ identity and may conflate percentage of place cells with the sensitivity and/or specificity of the subset of cells which are place cells (see Fig. 6D: increased PF sensitivity and specificity in *Pdzd8 KO* CA1PNs did not result in significantly increased spatial information – likely due to a modest reduction in place cell fraction). PF sensitivity and specificity were assessed separately (see below) to complement this measure. Spatial information was calculated as previously reported^27, 28^ using deconvolved events of active ROIs. Null distributions of 1000 scores were generated by circular position permutation as described above for spatial tuning curve calculations. Information scores were compared to null distributions to generate a p value and spatial information was reported as (1-p) such that higher values indicated greater spatial information content.

#### Place field properties

PF sensitivity was defined as the fraction of PF traversals during which an event was detected. PF specificity was defined as the fraction of all detected events occurring within PF bounds. For CA1PNs with multiple PFs, sensitivity and specificity were assessed collectively across PFs to yield one value per place cell. PF width was defined as the number of position bins separating PF bounds, multiplied by position bin length (2 cm).

### Within-dendrite analyses

For all within-dendrite analyses, which were motivated by potential differences in distributions of excitatory synaptic weights along single dendritic branches, apical trunks were excluded since they are largely devoid of spines^72^. For absolute measures of within-dendrite activity, including intradendritic activity variance and transient spread, data were normalized to mRuby3 intensity to control for differences in focality. Cells for which mRuby3 signal was not acquired were excluded from these analyses. mRuby3 signal was occasionally not acquired for cells with baseline jGCaMP7b signals that were clearly adequate on their own for motion correction since within-dendrite analysis was the initial intended purpose of a static red marker.

#### Activity variance

Intradendritic activity variance was calculated using the total counts of deconvolved events in an imaging session and the median mRuby3 intensity from the red channel motion-corrected time average for each subROI. To account for differences in the ranges of these values, coefficients of variation were used instead of absolute variance. To normalize against mRuby3 brightness, this intradendritic activity variance, *v*, was derived as the log_2_-normalized ratio of coefficients of variations for event counts and median brightness values which is equivalent to:

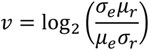

where *e* represents event counts and *r* represents mRuby3 median intensities for each subROI along an imaged dendritic ROI. Log_2_ normalization was chosen to generate values symmetric around zero; absolute fractions do not provide symmetrical measures and introduce biases with values greater than 2.

#### Transient spread

To calculate the spread of a given Ca^2+^ transient within an individual dendritic ROI, the dF/F_0_ signal from each corresponding subROI was indexed by the start and end frames of the whole-ROI transient and peak dF/F_0_ values were taken. To normalize these peak dF/F_0_ values to mRuby3 intensity, median mRuby3 intensities across subROIs were first normalized to their maximum value such that intensities then ranged between 0 and 1; otherwise, intensity values could be exceedingly high and squeeze out any differences in peak dF/F_0_ values. Next, peak dF/F_0_ for each subROI was divided by its own normalized mRuby3 intensity. Using these normalized peak dF/F_0_ subROI measurements, intradendritic transient spread, *s*, was calculated as:

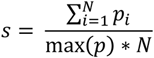

Where *p* is a vector of normalized peak dF/F_0_ values corresponding to subROIs of an individual dendrite for a given Ca^2+^ transient and *N* is the number of 2-µm subROIs from that dendrite. Thus, values fall on the interval [1/*N*, 1] where a value of 1 indicates that all subROIs showed equivalent peak dF/F_0_ during the transient. This measure naturally declines with longer dendritic ROIs (Fig. S9, B and C) but exceedingly long dendritic ROIs were relatively rare and robust differences between genotypes were observable out to 20 µm-long ROIs (Fig. S9, D and E).

#### Tuning curve correlations

To calculate intradendritic TC correlations for a given dendritic ROI, all possible combinations of subROI TCs were correlated with each other and correlation coefficients were binned by the anatomical distance separating the two subROIs. Mean correlation coefficients were then calculated for each distance bin to generate a within-dendrite TC correlation histogram. This process was repeated for all dendrites of a given genotype and compartment to calculate distributions of subROI TC correlations as a function of distance (Fig. 4, E and F).

### Detecting place field formation events

To detect *de novo* PF formation events, we began with mean PFs calculated from all laps^34^. However, in detecting the formative lap, we found previous methods to be susceptible to (1) missing PF formation events due to non-biological constraints on PF sensitivity, causing either no formation lap to be found or a lap to be chosen before which clear in-field activity was observable and (2) in our particular case, calling early formation laps without sufficient evidence that a PF was formed *de novo* as opposed to having existed prior to the imaging session. This is an issue when animals navigate environments which they have previously encountered, as was the case in this study.

To address these issues, we used an *N* = 2 Gaussian mixture model (GMM) to differentiate between “PRE” and “POST” laps relative to PF formation combined with post-hoc testing and specific conditions to minimize false positives. For each detected PF, a GMM was fit to a two-feature matrix with columns representing (1) a rolling average (weighted by Bartlett window) of within-field deconvolved events (“hits”) by lap and (2) lap numbers multiplied by exponentially decaying weights. Lap numbers were used to promote chronicity in PRE/POST lap classification; weighting lap numbers with an exponential decay was necessary to temper the influence of this feature. The GMM fit was randomly initiated 1,000 times for each PF and the best fit was taken. The PF was assumed to have existed prior to session start if (1) the model did not converge, (2) all laps were assigned to a single component (3) the mean hit/lap rate in PRE laps was greater than 1. If the model converged, then the first POST lap was used as the starting point of a 10-lap search window. The formation lap was determined to be the first lap within this window where the number of hits exceeded either the mean hit rate in POST laps or the mean hit rate of the search window; whichever was greatest. To account for bi-stable place cells in which activity toggles between two distinct PFs over laps, it was ensured (1) that the mean search window hit rate was greater than the mean PRE hit rate and (2) the lap with the most hits did not fall within the PRE component. To validate detected PF formation events, five conditions were imposed: (1) the same cell did not express an overlapping PF on the previous day, (2) the hits rates in PRE vs POST laps were significantly different by the Mann-Whitney *U* test, (3) the place cell fired within its PF on at least 1/5 laps after formation, (4) the place cell showed at least 35% specificity to its PF after formation, and (5) the PF did not form earlier than lap 10 to mitigate the possibility that it had formed prior to imaging.

### Analyzing place field formation events

#### Single-cell imaging data

For optogenetically-induced PFs in single cells, an ‘LED zone’ was defined to operationalize what activity should be considered induced. The LED zone was defined as a symmetric window, with a 15 cm radius, centered on LED onset position (Fig. 5C, green shaded area) on a cued, 2-meter treadmill belt. For each induction experiment, spatial tuning heatmaps were calculated as described above based on pre-induction (baseline), peri-induction, and post-induction laps using dF/F_0_. Baseline and post-induction heatmaps were flattened to generate tuning curves by taking the lap-averaged signal. Induction efficacy was calculated as the maximum difference between post-induction and baseline tuning curves within the LED zone. To account for potential differences in cell-to-cell excitability, as well as differences in jGCaMP7b plasmid expression that could squeeze the observable range of dF/F_0_, we normalized the baseline and post-induction tuning curves to the maximum observed LED response for a given ROI (Fig. 5C). An ROI was considered “induced” if its mean post-induction dF/F_0_ exceeded baseline dF/F_0_ in at least one position bin (50 4-cm bins were used) within the LED zone. This definition was designed to be inclusive rather than exclusive since cell bodies and dendrites showed variability in their degrees of optogenetically-induced spatial tuning that we wished to capture. Peak peri-formation velocity (figs. S4C; S10, D and F) was defined as the maximum instantaneous velocity observed from 10 cm preceding LED onset through to the animal’s position at LED offset (LED stimulation lasted 1 second.). Peak velocity was chosen as opposed to mean velocity because, occasionally, mice rested in some portion of this zone which drastically affected velocity measurements.

#### Population imaging data

For spontaneously-occurring PF formation events in population imaging data, peak peri-formation velocity was defined so as to approximate our measurement in optogenetic induction experiments. Peak velocity was taken from a window starting 10 cm before the first boundary of the detected PF and ending with the PF end boundary. Widths of *de novo* PFs were calculated as an average over all laps, i.e. agnostic to when the PF formed.

### Statistical analysis

For all group comparisons, normality was assessed using the Kolmogorov-Smirnoff test. When one or more groups significantly diverged from a normal distribution, the nonparametric, two-tailed Mann-Whitney *U* test was used. Otherwise, two-sided *t* tests with Welch’s correction for heteroscedasticity were used. Descriptive statistics for non-normal distributions were reported as median ± inter-quartile range; normal data were reported as mean ± standard error of the mean. For multi-way comparisons, ANOVA was typically used except in the case of missing values in which case a Mixed Effects model was used with the Geisser-Greenhouse correction for asphericity. For single-cell imaging data (Figs. 2-5), unless otherwise noted, Ns represent cell-sessions (N cells x N days imaged) since dendritic ROIs varied day-by-day both in location along the dendritic arbor (Fig. S5) and in spatial tuning properties (Fig. 1F).

---

**Fig. S1.**
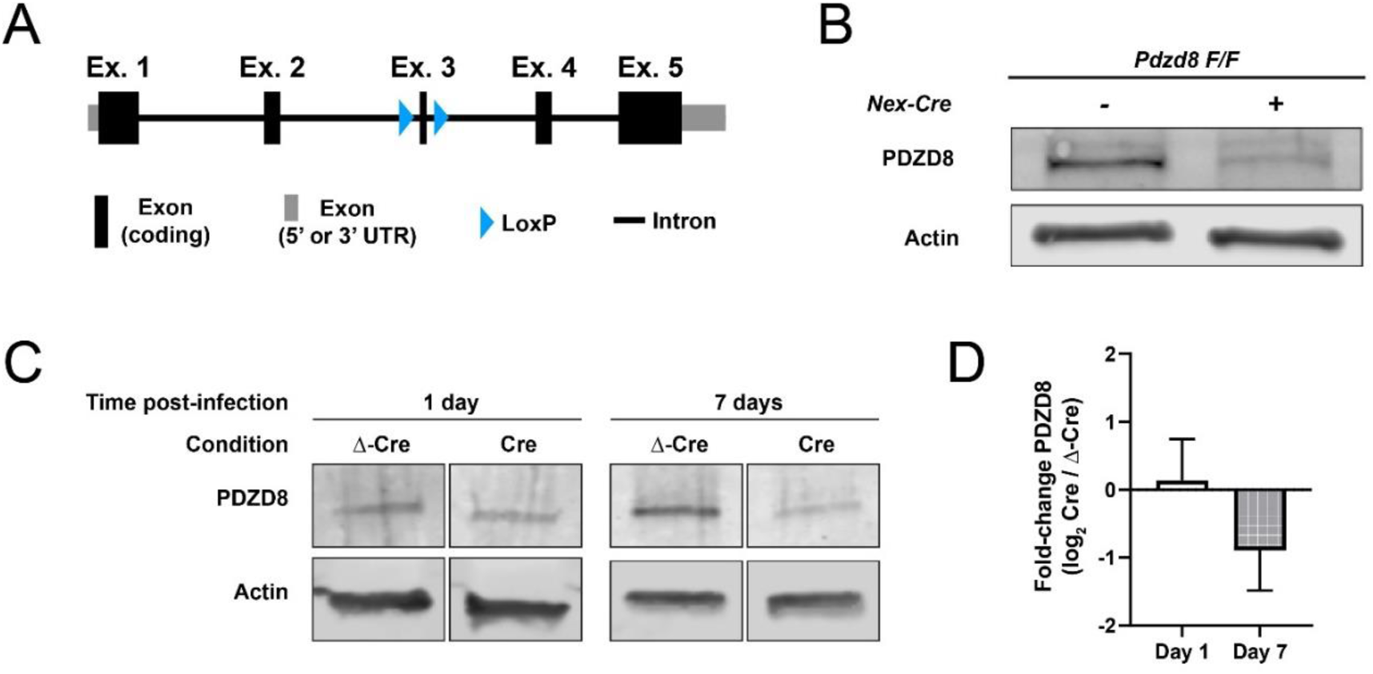
*Pdzd8 F/F* conditional knockout mouse line. (**A**) Diagram showing conditional knockout strategy for *Pdzd8*. Exon 3 is surrounded by loxP sites. Cre-mediated recombination excises the encircled DNA, creating a frame-shift mutation which results in total knockout of the *Pdzd8* gene. (**B**) Western blot of brain tissue from *Pdzd8 F/F; Nex− Cre −/−* and *+/−* mice in which *Pdzd8* is knocked out from glutamatergic neurons of the dorsal telencephalon^73^. Samples from Cre-positive mice show loss of PDZD8 protein but not Actin control. (**C**) Example western blots of primary cultured neurons from *Pdzd8 F/F* embryos showing PDZD8 levels at 1 and 7 days following infection with a lentiviral construct expressing either Cre or inactive Δ-Cre control (see Methods). (**D**) Quantification of PDZD8 signal fold-change. At 7 days post-injection, the earliest that CA1PNs were imaged *in vivo* following single-cell electroporation (Fig. 1A), PDZD8 protein is reduced. PDZD8 signal was normalized to total protein for quantification.

**Fig. S2.**
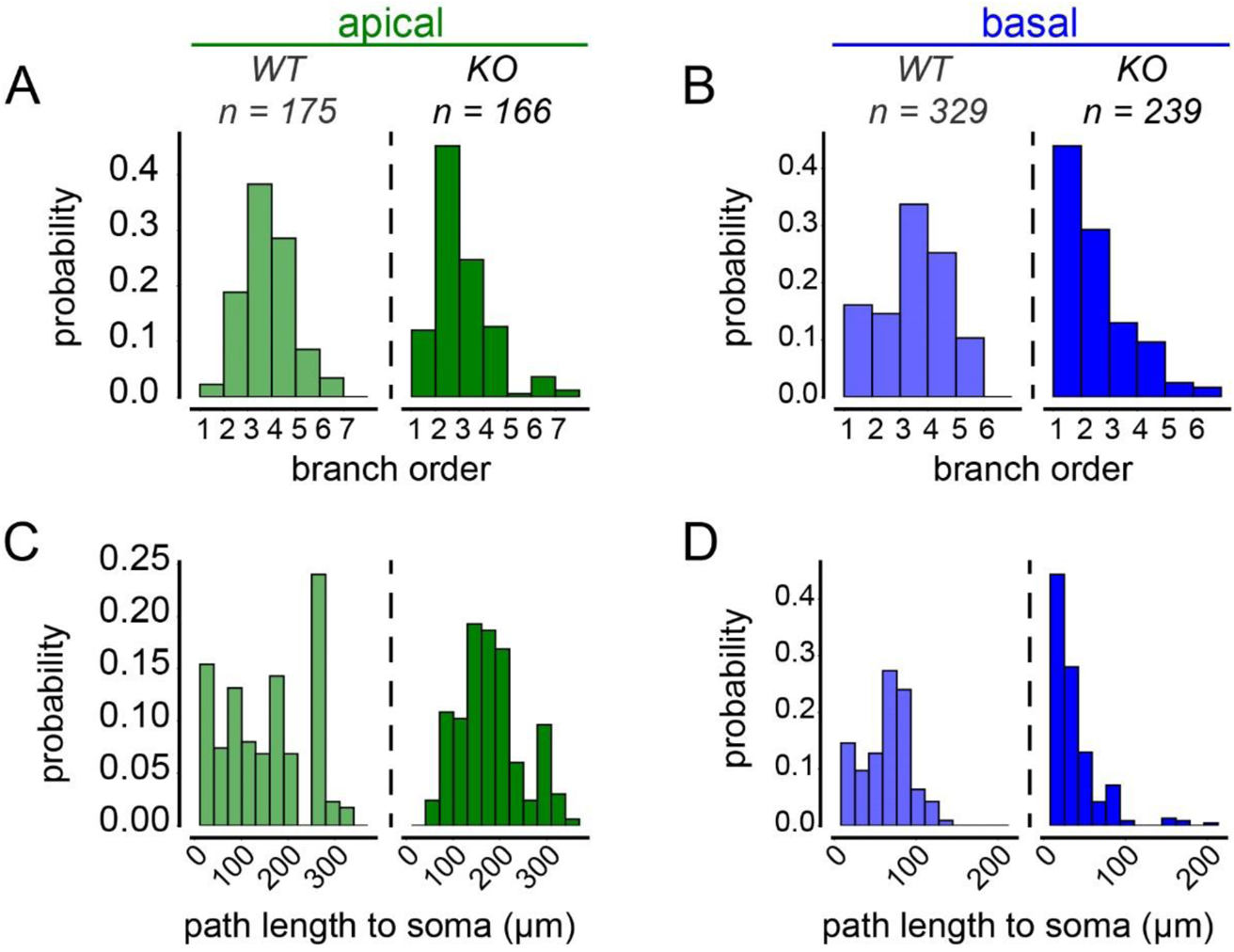
Morphological properties of imaged dendrites. (**A, B**) Distributions of branch orders for imaged apical (green, A) and basal (blue, B) dendrites of *Pdzd8 WT* (transparent) and *KO* (opaque) CA1PNs. (**C, D**) Path lengths from imaged dendrites to respective soma, as shown in (A, B).

**Fig. S3.**
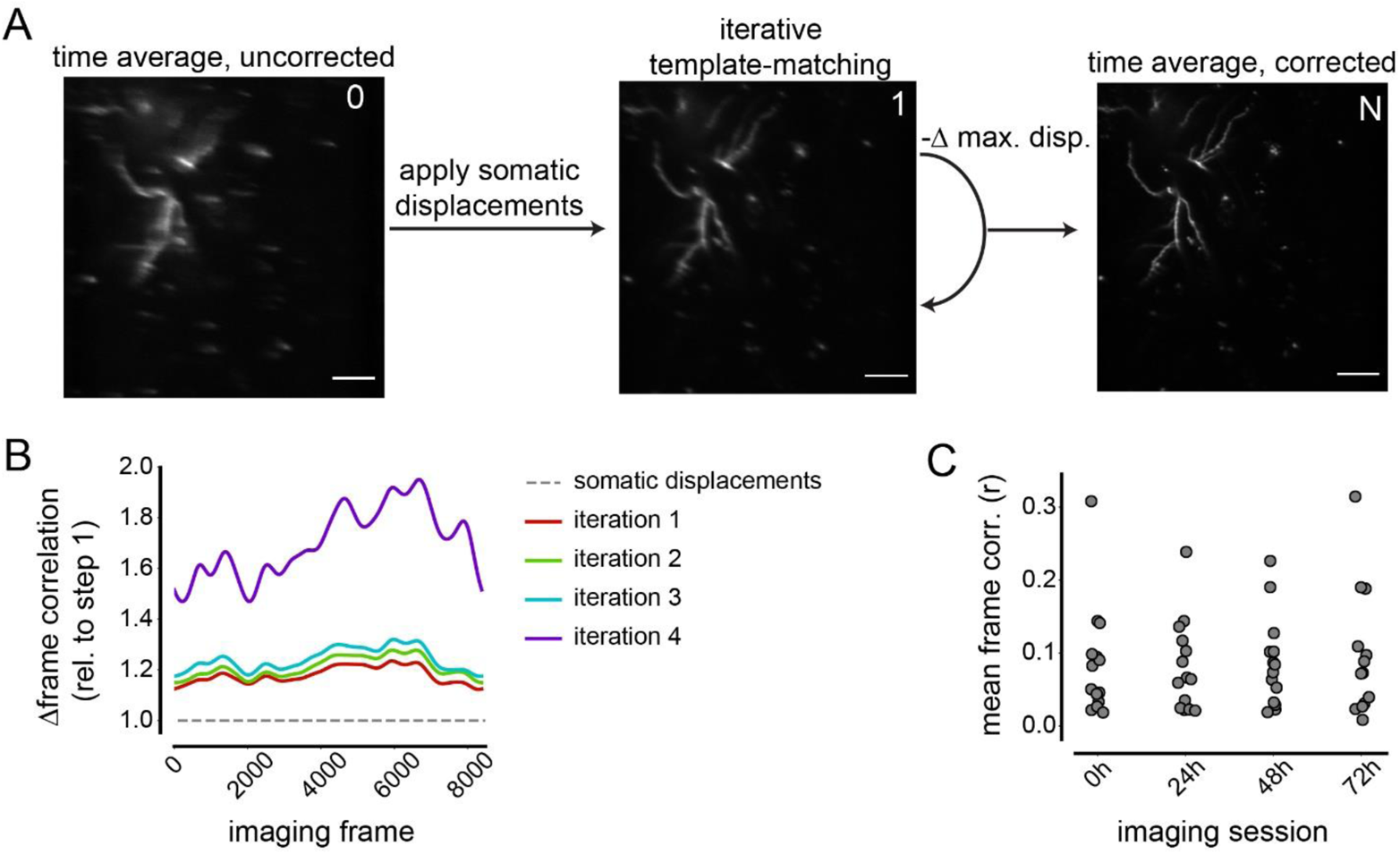
Iterative template-based motion correction in sparse dendritic fields of view. (**A**) Schematic showing motion correction for example apical dendritic field of view (FoV) from a single-cell, multi-plane imaging session. Time-averaged FoVs are shown before motion correction (left), after applying somatic displacements to dendritic data to generate a first estimate (middle), and after iterative correction steps using the previous motion-corrected dendritic time average as a template (right) with increasingly strict constraints on maximum allowable displacements (-Δ max. disp.). Signals shown are from bReaChes-mRuby3 fusion protein co-expressed with jGCaMP7b. Scale bar 30 µm. Contrast adjusted to aid visualization. (**B**) Improvements in frame-to-time average correlation with successive rounds of iterative template-based motion correction for example FoV shown in (A). Values are normalized to initial starting point generated by application of somatic displacements (dashed grey line). Correlations were smoothed with a 100-frame window before calculating improvements by iteration. (**C**) Mean frame-to-time average correlations for each imaged dendritic focal plane plotted by imaging day. Individual frames typically contained little signal and thus weakly correlated with time-averaged FoVs, hence the need for strategy depicted in (A). Pearson correlation coefficients were used.

**Fig. S4.**
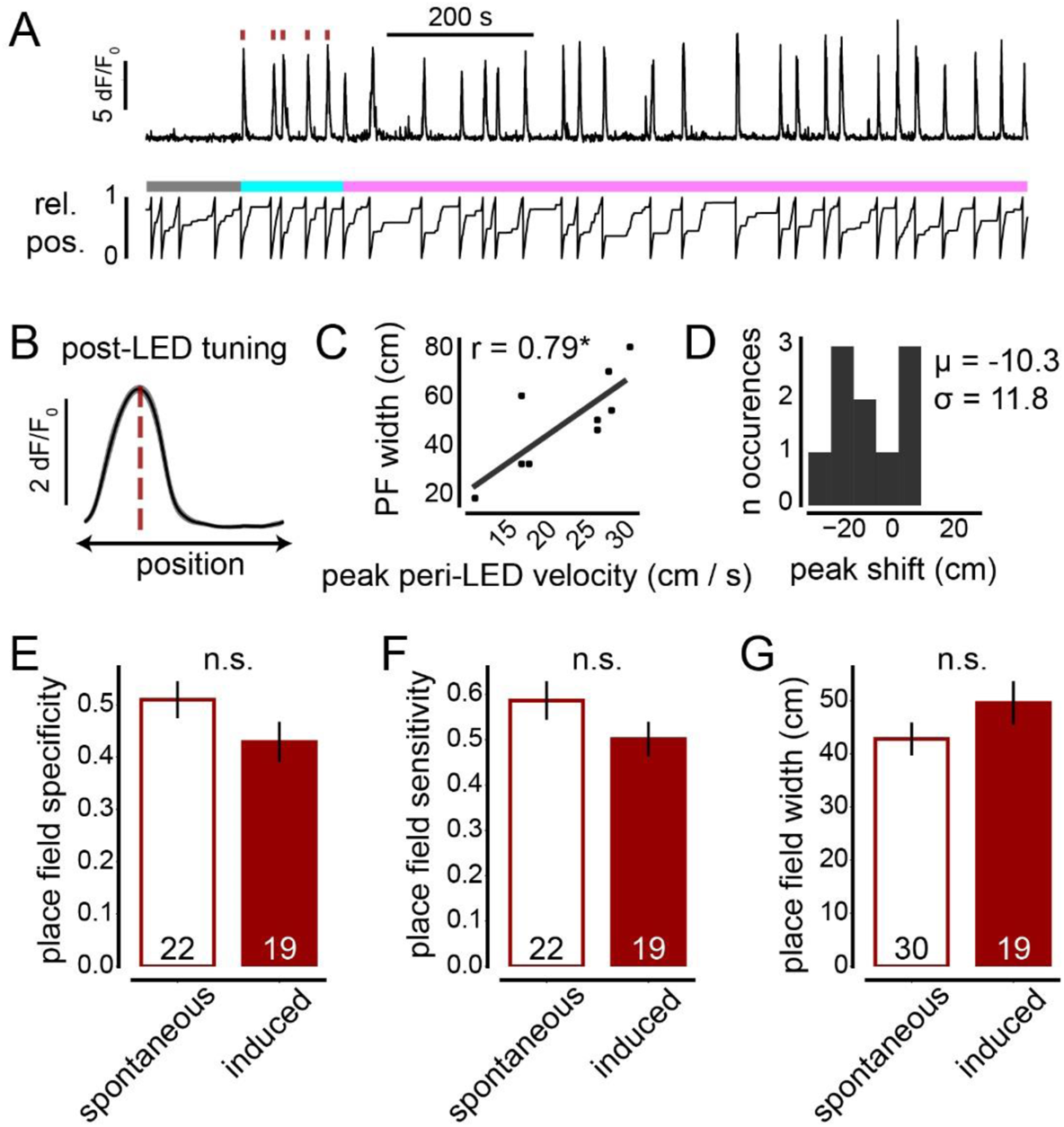
Optogenetically induced place fields show evidence of behavioral timescale synaptic plasticity and resemble naturally-occurring, spontaneous place fields. (**A**) Example somatic jGCaMP7b trace of place field (PF) induction experiment (top). Soma shows no discernible activity in pre-LED (baseline) period, reliable responses to LED stimulation (red ticks), and consistent subsequent activity. Activity is evaluated with respect to animal’s relative position (rel. pos.) along treadmill belt (bottom) to evaluate spatial tuning. Baseline (grey), induction (cyan), and post-induction (magenta) laps are indicated by color bar (see Fig. 1, D-F). (**B**) Post-induction spatial tuning from example in (A), shown as mean dF/F_0_ by position bin (50 bins from 2-meter track, averaged over 24 post-induction laps). (**C**) Relationship between animal velocity around time of first LED exposure and width of resultant PF in *WT* CA1PNs, as previously observed in plateau potential-dependent behavioral time scale synaptic plasticity (BTSP)^31^. p < 0.05*, Pearson correlation analysis. N = 9 induced place fields from 9 cells. (**D**) Histogram showing peak locations of induced *WT* PFs shown in (C), as assessed on post-induction laps, relative to LED onset. Negative peak shifts result from asymmetric plasticity kernel underlying BTSP^31^. Mean (µ) and SD (σ) are shown on plot. (**E-G**) Comparison of spontaneous and induced somatic PF properties. PFs overlapping with a nominal “LED zone” (see Fig. 5C, Methods) were considered to be induced. Data are shown as mean ± SEM. Two-sided unpaired t-tests were used.

**Fig. S5.**
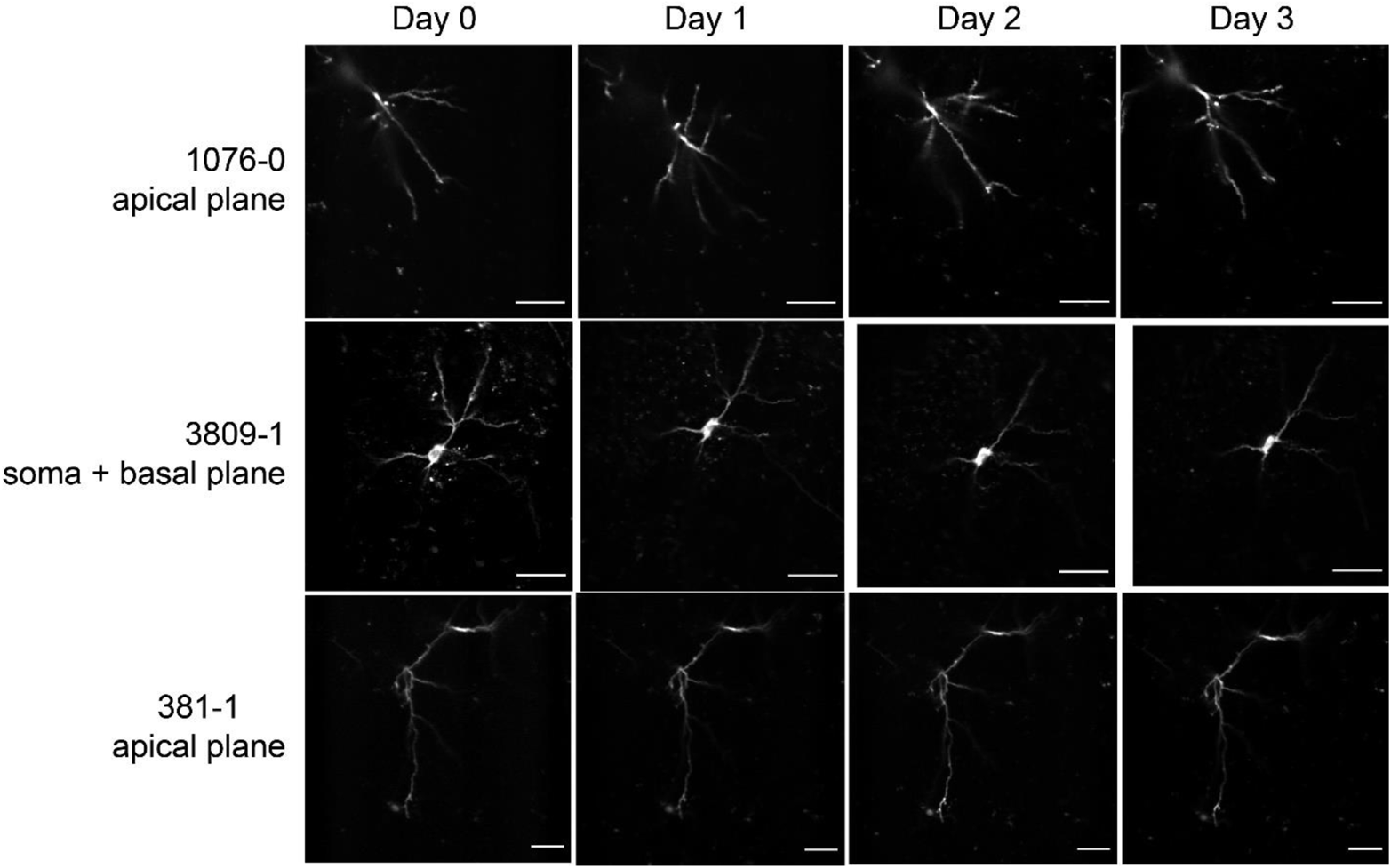
Mixed success in precisely tracking CA1PN dendrites across days. Motion-corrected time-averaged fields of view showing static mRuby3 signal from each single-cell imaging session across all days (columns) from 3 example cells (rows). One of two imaged planes is shown for each cell. Top two rows (1076-0, 3809-1) show example cells for which identical planes were not acquired across days. Bottom row shows example cell with consistent apical dendritic focal planes across days. Contrast was adjusted for ease of comparison. Scale bar 40 µm.

**Fig. S6.**
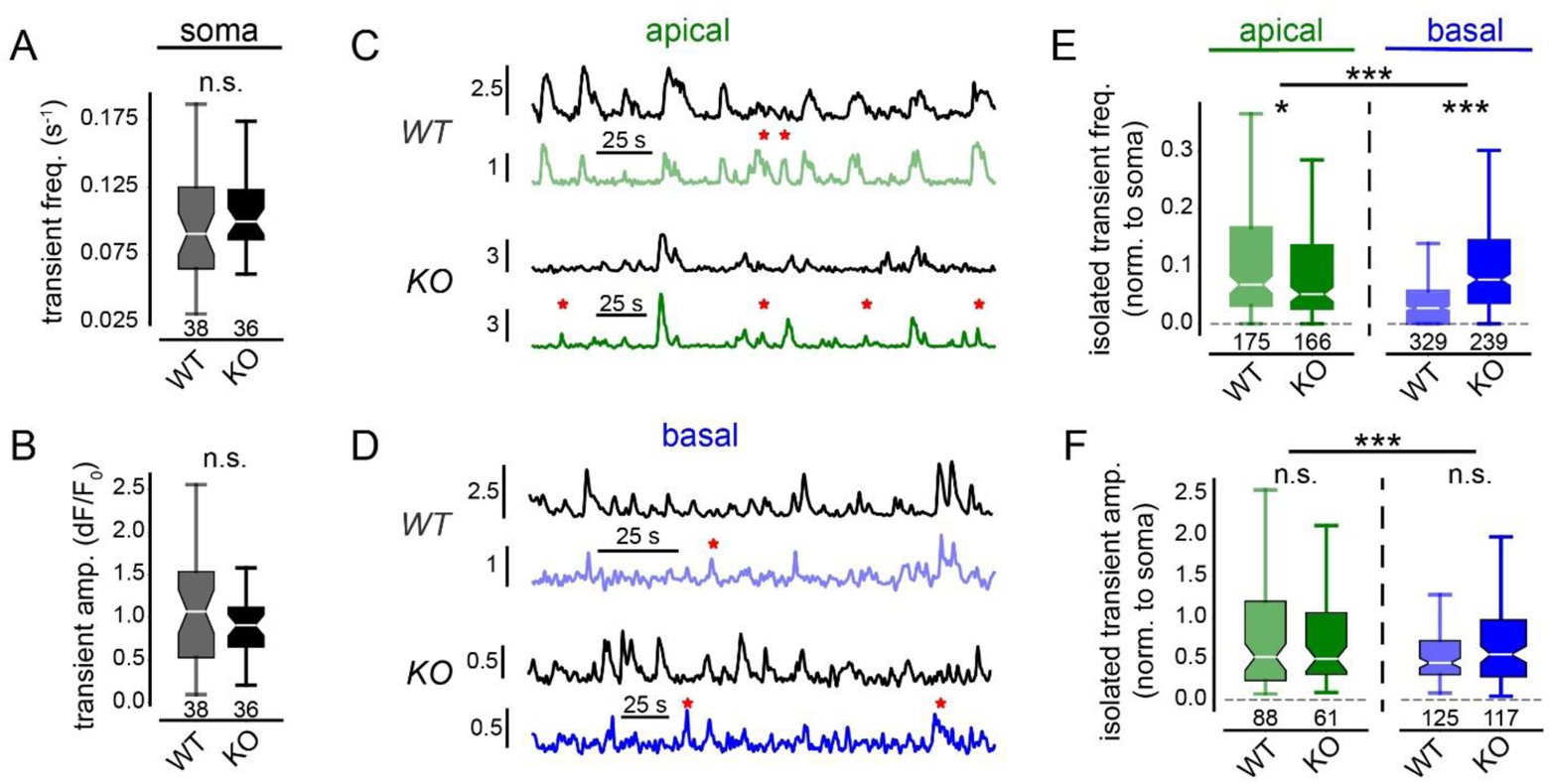
Event-based comparison of Ca^2+^ dynamics in *Pdzd8 WT* and *KO* soma and dendrites. (**A**) Somatic Ca^2+^ transient frequency (*WT*: 0.090 ± 0.61, n = 38; *KO*: 0.100 ± 0.037, n = 36; p > 0.05). (**B**) Mean amplitude of somatic Ca^2+^ transients (*WT*: 1.09 ± 0.13, n = 38; *KO*: 0.94 ± 0.074, n = 36; p > 0.05). (**C**) Example traces showing isolated dendritic Ca^2+^ transients in *WT* (transparent) and *KO* (opaque) apical dendrites relative to co-acquired somatic traces from the same cell. Red asterisks indicate Ca^2+^ transients identified as isolated, i.e. not overlapping in time with a somatic transient. Vertical scale bars indicate dF/F_0_. (**D**) Example traces showing isolated basal dendritic Ca^2+^ transients as in (C). (**E-F**) Mean frequency (E) and amplitude (F) of isolated Ca^2+^ transients detected in apical and basal dendrites of *Pdzd8 WT* and *KO* CA1PNs. Dendritic values were normalized to mean somatic values (see Methods). 2-way ANOVA of frequency with post-hoc t-tests, compartment effect: F_1,907_ = 21.61, p < 0.001; genotype effect: F_1,907_ = 9.05, p < 0.01; interaction: F_1,907_ = 38.69, p < 0.001. 2-way ANOVA of amplitude, compartment effect: F_1,389_ = 7.07, p < 0.01; genotype effect: F_1,389_ = 2.47, p > 0.05; interaction: F_1,389_ = 0.00058, p > 0.05. LED-evoked events were excluded from all analyses. Ns shown in (E) represent total dendrites imaged across days since frequency could be measured in ROIs that showed no isolated events. Ns shown in (F) represent total dendrites across days that fired at least one isolated transient.

**Fig. S7.**
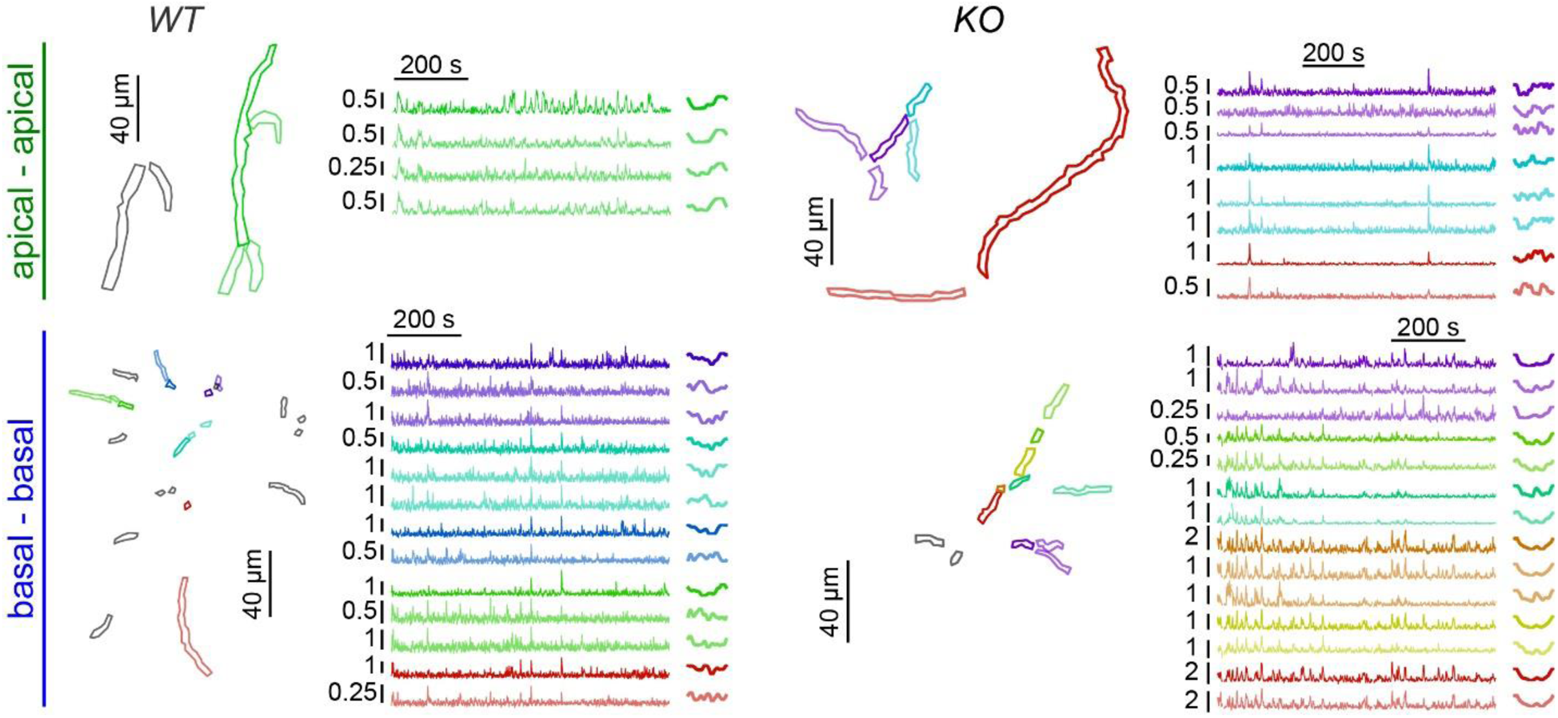
Analysis of connected, co-imaged dendrites. For apical-apical (top row) and basal-basal (bottom row) dendrite pairs from *Pdzd8 WT* (left column) and *KO* (right column) CA1PNs, co-imaged ROIs (left) are plotted along with tuning curves (middle) and dF/F_0_ traces (right) of connected dendrites. Imaged dendrites for which no “parent” ROI was imaged are drawn in gray. For connected dendrites, the “parent” dendrite ROI and trace (closest to soma by branch order) are drawn (opaque) along with those of their directly-connected “child” dendrites (same color, decreased opacity). Sets of connected dendrites are separated by individual opaque/transparent color schemes. For child dendrites which also act as parent dendrites (see cyan and violet traces in top right quadrant), the parent color takes precedence for ROI plotting and the corresponding TC and trace are plotted separately for each group.

**Fig. S8.**
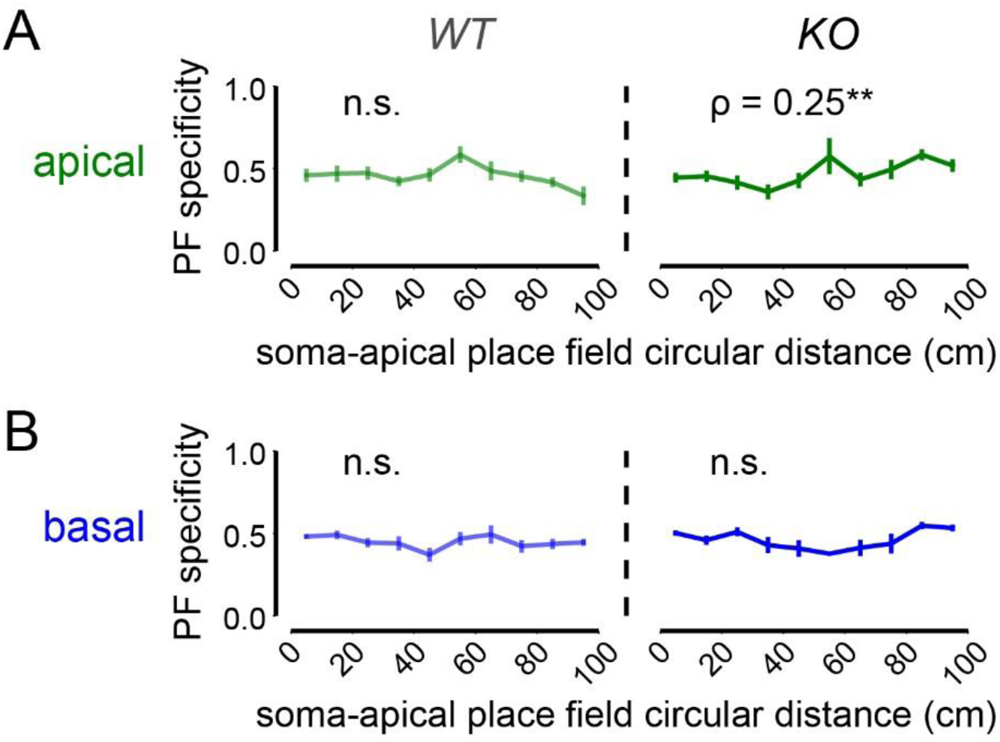
Relationship between soma-dendrite place field co-tuning and place field quality. (**A**) Place field (PF) specificity plotted as a function of soma-apical PF circular distance binned every 20 cm for *Pdzd8 WT* (transparent) and *KO* (opaque) CA1PNs. Spearman correlation analysis was used on non-binned data. N = 145 *WT* and 164 *KO* PFs from apical dendrites belonging to place cells. (**B**) PF specificity and soma-dendrite PF circular distances for basal dendrites as in (A). N = 318 *WT* and 302 *KO* PFs. Error bars represent SEM of binned values.

**Fig. S9.**
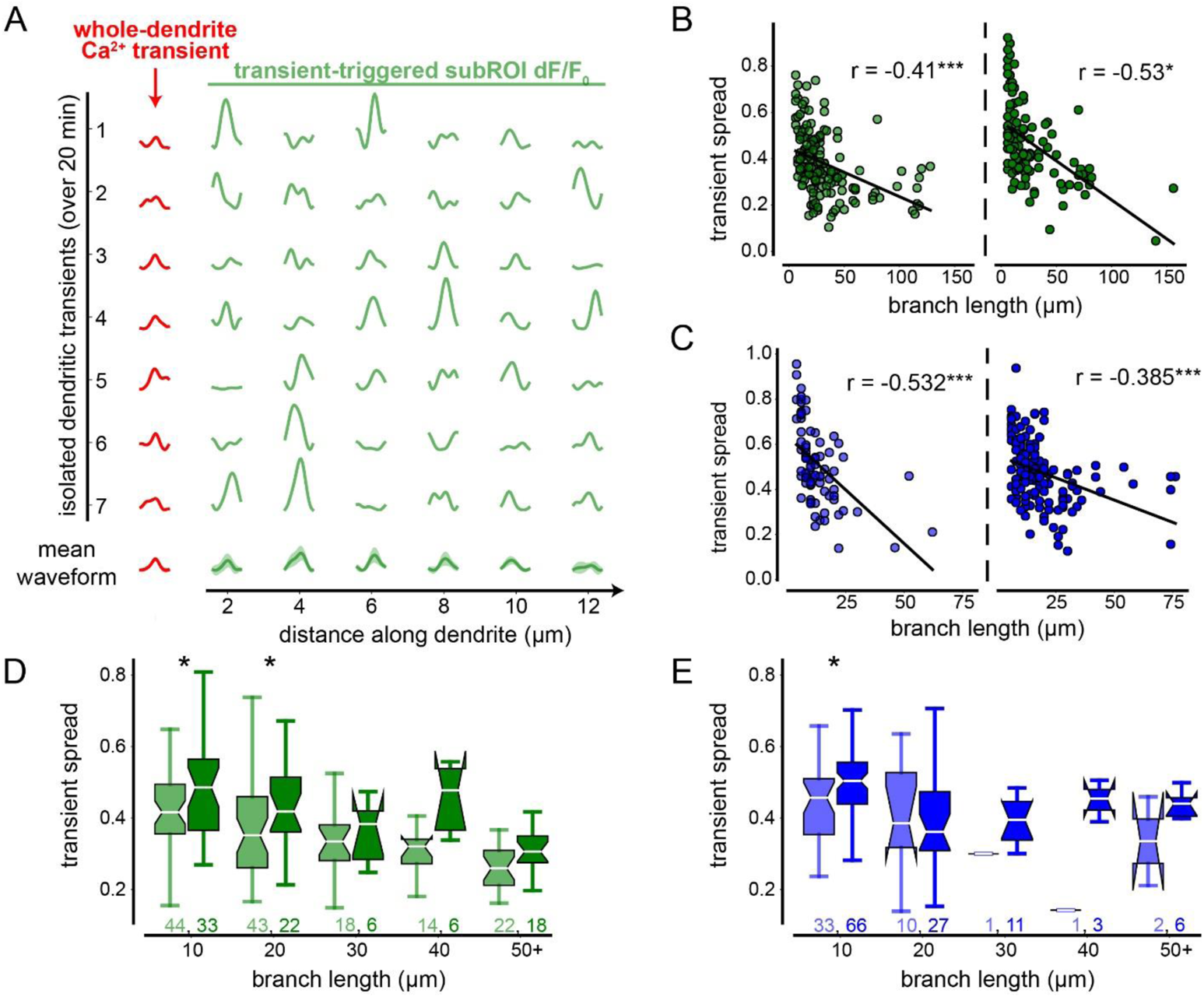
Isolated dendritic Ca^2+^ transients: variable points of initiation and branch-length-dependence of spread. (**A**) Isolated Ca^2+^ transients from an example apical dendritic ROI. Each row represents a successive isolated transient recorded over a 20-min. imaging session. Red waveforms at left represent overall waveform from entire ROI, centered on transient peak amplitude. Green waveforms at right represent corresponding subROI dF/F_0_ from same imaging frames. subROI traces are plotted in anatomical order along whole ROI. Mean waveforms for whole ROI (red) and each subROIs (green) across all isolated transients are plotted at bottom with shaded SEM. Variable locations of maximal amplitude (i.e. potential point of initiation) across events indicates that within-dendrite dynamics are input-driven. (**B**) Intradendritic transient spread of *Pdzd8 WT* (transparent, left) and *KO* (opaque, right) apical dendrites plotted as a function of branch length. subROI dF/F_0_ amplitudes are normalized to mRuby3 intensity to account for differences in focality. Black lines represent linear regression fit. Pearson correlation coefficients shown on plots. Here the full domain is included to illustrate that transient spread obligatorily decreases with segment length (see Methods). (**C**) Basal intradendritic transient spread as shown in (B). (**D**) Spread of isolated transients in *Pdzd8 WT* (transparent) and *KO* (opaque) apical dendrites, binned by branch length to account for trends shown in (B, C). Boxes range from lower to upper quartiles with line at median; whiskers show range of data within 1.5 * (Q3 - Q1) of box boundaries. (**E**) Spread of isolated transients in basal dendrites as in (D). 2-way ANOVA indicates interaction between genotype and branch length (F_1, 25_ = 4.81, p < 0.001). *p < 0.05, ***p < 0.001.

**Fig. S10.**
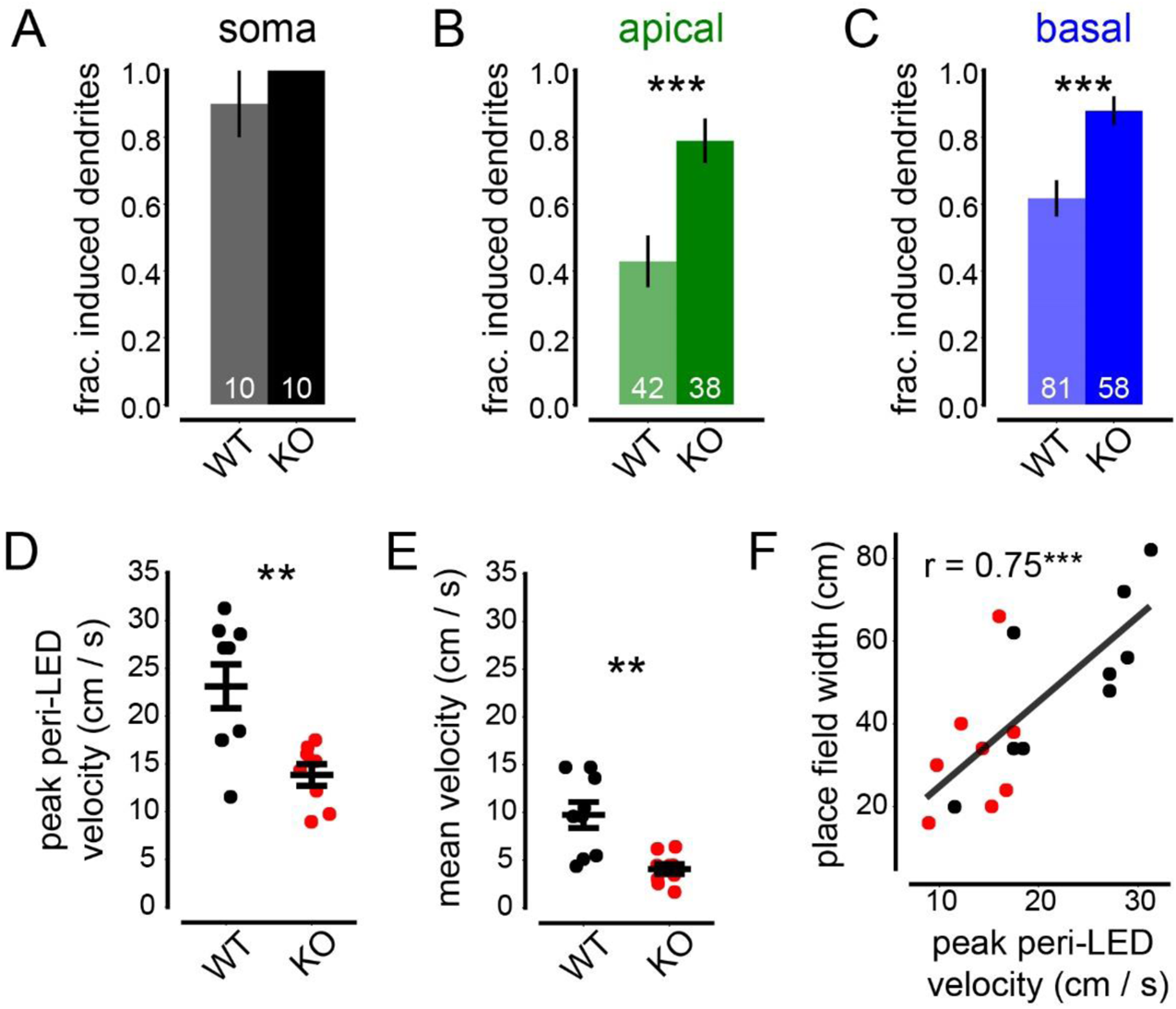
Place field induction success rate and animal behavior. (**A-C**) Fraction successfully induced somatic (A), apical (B), and basal (C) ROIs by genotype. Induction success criterion did not require a significant place field; only a post-stimulation increase in activity within the nominal “LED zone” relative to baseline (see Fig. 5C, Methods). (**D**) Peak velocity near LED zone on the first lap of optogenetic stimulation during induction sessions. (**E**) Mean velocity across the entire induction session. (**F**) Width of induced somatic place fields, as calculated from post-stimulation laps on induction day, plotted as a function of peak peri-LED velocity as shown in (D). *Pdzd8 WT* (black) and *KO* (red) data are shown together. Black line denotes linear regression fit. Pearson correlation coefficient is shown. Distributions were compared using two-sided unpaired t-tests and Mann-Whitney U tests. Error bars represent SEM. **p < 0.01, ***p < 0.001.

**Fig. S11.**
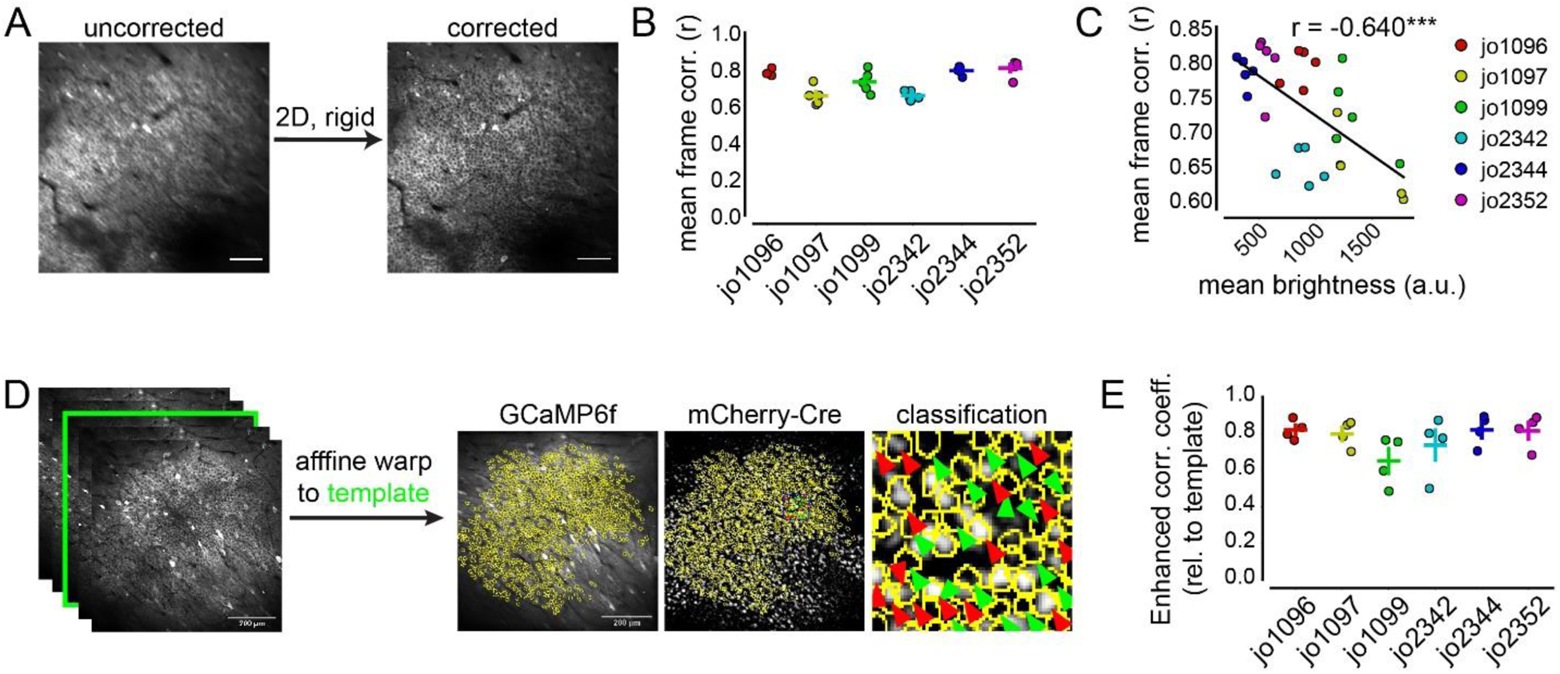
Motion correction and multi-day cross-registration of mixed population Ca^2+^ imaging data. (**A**) Example field of view (FoV) time averages of GCaMP6f signal before and after motion correction using rigid 2-dimensional translation approach. Scale bar 100 µm. (**B**) Mean frame-by-frame correlations with time-averaged FoV. Each mouse shown in separate color, individual points represent imaging sessions. Crosses indicate mean ± SEM. (**C**) Relationship between mean frame-by-frame correlation and mean overall FoV brightness. Imaging sessions are color-coded by mouse ID. Black line denotes linear regression fit. Pearson correlation coefficient shown on plot (***p < 0.001). (**D**) Schematic showing affine transformation-based approach to align all imaging sessions from a particular animal to a chosen “template” day (see Methods). Scale bar 200 µm. Regions of interest (ROIs) are drawn over aligned, concatenated dataset using GCaMP6f time average. Co-aligned mCherry-Cre signal is used to classify cells as *Pdzd8 WT* (mCherry (-), green arrows) or *KO* (mCherry (+), red arrows). Classification example is magnified from blue box in mCherry-Cre FoV. (**E**) “Enhanced correlation coefficient”^66^ quantifying goodness of fit between the time average of each imaging session and that of its corresponding template session (template-template coefficients are omitted). Data organized my mouse ID as in (B, C).

**Fig. S12.**
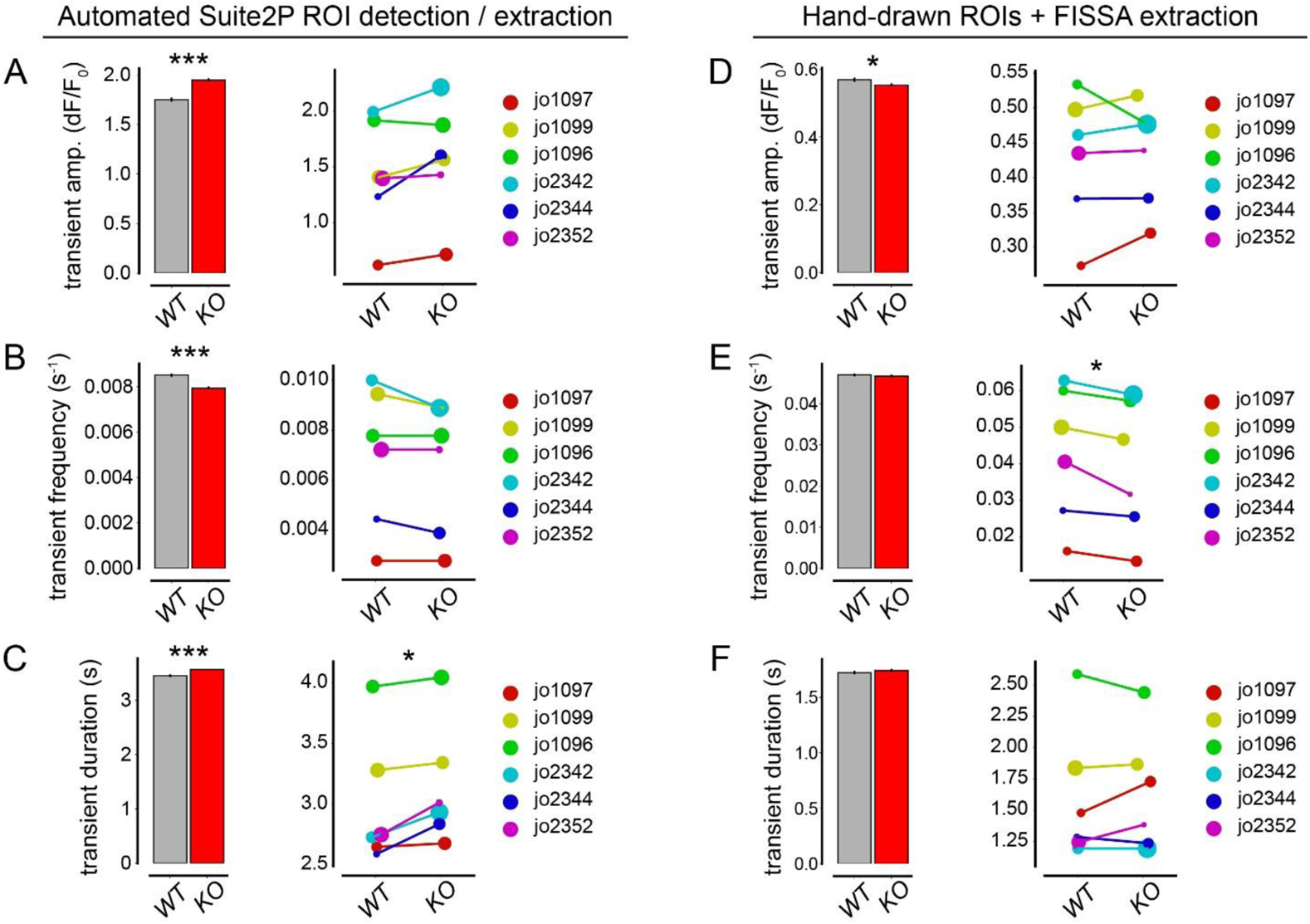
Ca2+ transient properties of Pdzd8 WT and KO CA1PNs. (**A**) Left: Bar plot comparing Ca^2+^ transient amplitudes (mean value across all 5 days for each ROI) for *Pdzd8 WT* (grey, n = 2,719 cells from 6 mice) and *KO* (red, n = 3,260 cells from same 6 mice) CA1PNs. Right: Mean values plotted by animal. Dot sizes are weighted by number of ROIs relative to total ROIs within-animal (e.g. *N_WT_* / (*N_WT + KO_*) to illustrate balance of *Pdzd8 WT* vs *KO* CA1PNs in each subject. (**B-C**): Ca^2+^ transient frequency (B) and duration (C) shown as in (A). (**D-F**): Ca^2+^ transient properties shown as in (A-C) extracted from hand-drawn ROIs using FISSA^70^ to decontaminate neuropil signals as similarly done by Suite2P package^67^. Hand-drawn ROIs were explored in case automated, activity-based Suite2P ROI segmentation excluded weakly active or silent cells. Paired t-tests were used for within-animal comparisons. Across-animal distributions were compared using Mann-Whitney U tests. *p < 0.05, **p < 0.01, ***p < 0.001. Error bars represent mean ± SEM.

**Table. S1.** Single-cell imaging Ns by genotype, ROI type, and day.

|  | WT |  |  | KO |  |  |
| --- | --- | --- | --- | --- | --- | --- |
| Day | soma | apical | basal | soma | apical | basal |
| 1 (0h) | 10 | 42 | 81 | 9 | 38 | 58 |
| 2 (24h) | 10 | 41 | 88 | 9 | 43 | 58 |
| 3 (48h) | 10 | 45 | 80 | 9 | 45 | 58 |
| 4 (72h) | 8 | 47 | 80 | 9 | 40 | 65 |

**Movie S1.** Somatic (left) and apical dendritic (right) imaging planes of example cell shown in Fig. 1B (right) shown side-by-side during a place field induction lap. jGCaMP7b response to a single, 1-second LED pulse is shown. Frames are motion-corrected and shown at actual speed. Data acquired with 40X objective at 1.0X optical zoom.

**Movie S2.** Population GCaMP6f dynamics in CA1 pyramidal layer during random foraging spatial navigation as in Fig. 6. Frames are motion-corrected and shown at actual speed. Data acquired with 16X objective at 1.2X optical zoom.

